# Understanding flux switching in metabolic networks through an analysis of synthetic lethals

**DOI:** 10.1101/2024.03.05.583461

**Authors:** Sowmya Manojna, Tanisha Malpani, Omkar S. Mohite, Saketha Nath, Karthik Raman

**Author notes:** These authors contributed equally to this work (Sowmya Manojna, Tanisha Malpani).

## Abstract

Biological systems are extremely robust and exhibit high levels of redundancy for multiple cellular functions. Some of this redundancy manifests as alternative pathways in metabolism. Synthetic double lethals in metabolic networks comprise pairs of reactions, which, when deleted simultaneously, abrogate cell growth. However, when one reaction from such pairs is removed, the cell reroutes its metabolites through alternative pathways. Very little is known about the set of reactions through which fluxes are rerouted. Analysing this redistribution would help us to uncover the linkage between the reactions in a synthetic double lethal and also understand the complexity underlying the reroutings. Studying synthetic lethality in the context of pathogenic bacteria can offer valuable insights into therapeutic interventions. In this work, we propose a constraint-based approach to unravel these alternate pathways and complex interdependencies within and across metabolic modules. The approach involves a generic optimisation that minimises the extent of rerouting between two reaction deletions, corresponding to synthetic lethal pairs. We also include a systematic analysis of synthetic lethals by identifying the reaction classes that make up these synthetic lethals. We applied our computational workflow to several existing high-quality genome-scale models to show that these rerouted reactions span across metabolic modules, thereby illustrating the complexity and uniqueness of metabolism. Our results provide interesting insights into the organisation of metabolic networks and their redundancy.

The algorithm is available at https://github.com/RamanLab/minRerouting.

## 1 Introduction

Robustness in the face of environmental perturbations is an essential attribute of microorganisms [1–3]. This robustness is often achieved by the presence of multiple alternate pathways that achieve similar metabolic functions [4, 5]. The redundancy introduced by alternate pathways comprises a large fraction of most metabolic networks [6–8]. The redundancies in reaction pathways also show surprising variance in their distribution [9, 10]. While some of the alternate pathways are very simple, arising due to gene duplication, other alternate pathways could be extremely complex, with compensating reactions spanning different metabolic subsystems [9, 10].

A straightforward method of studying these alternate pathways involves the identification of *synthetic lethals* in a metabolic model [10]. Synthetic lethals are sets of genes/reactions where only the simultaneous loss of all genes/reactions in the set leads to abrogation of cell growth [11]. When only one of the reactions is deleted, the cell is able to summon alternate pathways to ensure its survival. In many cases, this is made possible through a complex rerouting of fluxes in the metabolic network which exploits the redundancy in metabolism. However, very little is known about how these organisms reroute their fluxes, and how various reactions in the cell can compensate for one another. Previous uses of genome-scale models, including GIMME, iMAT, RELATCH, for the study of flux distributions have focused on a given condition, the final steady state of the cell, without considering the prior reference state of the organism [12–14]. REMI and deltaFBA are algorithms that integrate differential expression of transcriptome and metabolome with the flux distributions between two different states, a WT state and a mutant state [15, 16]. These algorithms require gene expression data, which is only sometimes available. Considering cells have a high order of redundancy and synthetic lethals, we require a method that can computationally predict the rewiring of metabolism. Previous studies [17, 18] used FBA to study redundancy using synthetic lethals in *E. coli* and other bacteria. They used FBA directly to optimise the single deletions for the biomass objective and find the differences in the flux distributions. However, this can ignore the biological costs associated with altering flux in an organism, which may result in sub-optimal biomass production.

To identify this set of reactions that come into effect to rescue the cell from a non-lethal deletion, we propose a novel approach termed *minRerouting*. By solving a minimum *p*-norm problem, *minRerouting* can simultaneously solve for flux distributions that satisfy the stoichiometric constraints, maximise the biomass constraint (with a slack, *γ*), and also minimise the number of reactions with varying metabolic flux values. It gives us the set of reactions with an altered flux as the output and, thus, helps us understand the redundancies that help make the wild-type organism robust against perturbations and mutations. In addition to studying synthetic lethals and redundancy, this approach is also ideal for exploring and understanding indispensable changes in cellular metabolism between two different conditions, for instance, a healthy state and a diseased state.

Robustness arising from double lethals has been studied previously. It has been proposed that double lethal pair robustness in an organism can be ascribed to two classes of reaction pairs—Plastic Synthetic Lethals (PSL) and Redundant Synthetic Lethals (RSL) [18]. PSL pairs are reaction pairs where only one reaction is active, while the other reaction is inactive. The second reaction becomes active only when the first reaction is inactive. RSL pairs are reaction pairs where both the reactions are active simultaneously; yet, the loss of one does not abrogate growth. It has also been shown that these classes are conserved even across different nutrient conditions. The very presence of two distinct reaction pair classes calls for us to analyse the cause behind such selective activation. Are the inactive reactions more “metabolically costly” than the active ones? What kind of reactions make up the RSL pairs, especially when they are both simultaneously active?

To answer these questions and explore the structure of metabolic networks and their underlying redundancy, we also make use of parsimonious Flux Balance Analysis (pFBA; [19]). In pFBA, the genes and reactions in a metabolic network are classified based on the maximum and minimum flux associated with them. We obtain the classes of each double lethal pair using a novel workflow of Flux Variability Analysis, analyse the reaction types contributing to the PSL and RSL classes, and uncover interesting patterns in their distribution.

## 2 Methods

### 2.1 Flux Balance Analysis

Flux Balance Analysis (FBA) [20, 21] is a constraint-based approach that is used to predict the steady-state flux distribution in a given organism’s metabolic network. FBA employs a Linear Programming (LP) formulation, with an objective to maximise the biomass flux under certain flux and stoichiometric constraints. The formulation of FBA is as follows:

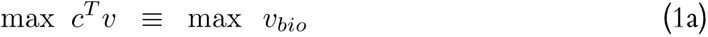

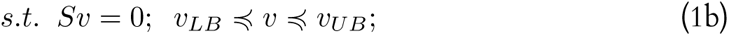

Here, *v* represents the flux vector and the *j^th^* entry corresponds to the flux through the *j^th^* reaction, *c* represents the objective function. Typically *c^T^ v* = *v_bio_*, where *v_bio_* is the biomass flux, *S* represents the stoichiometric matrix of dimensions *m × r*, where *m* is the number of metabolites and *r* is the number of reactions, and *v_LB_* and *v_UB_* represent the permissible lower and upper bounds of the reaction fluxes. FBA has been experimentally validated in many scenarios [22, 23], and has widespread applications [24]. An important extension of FBA is MoMA (Minimisation of Metabolic Adjustment; [25]), which seeks to identify a minimally different flux from the wild-type flux (by minimising the *l*_2_ norm of this difference), that is compliant with the new constraints imposed by a perturbation such as a reaction deletion.

### 2.2 Identification of Synthetic Lethals

Fast-SL [26, 27] is an efficient algorithm that identifies synthetic lethals by systematic pruning of the search space and exhaustive enumeration from the remaining reactions. Fast-SL rapidly identifies synthetic lethals and scales well for higher-order lethals. In this paper, we used Fast-SL to identify synthetic lethal reaction pairs in a given genome-scale metabolic model. We used a threshold of 10*^−^*^5^ to identify active reactions in Fast-SL.

### 2.3 minRerouting Formulation

We define the ‘minRerouting set’ as the minimal reaction set comprising all the reactions that have a modified flux following the individual deletion of synthetic double lethal reactions. This can be seen in context in Figure 1, which represents a toy metabolic network comprising nine reactions and eight metabolites. The metabolites *A* and *B* are the ‘input’ metabolites and *G* is the ‘output’ metabolite.

**Fig. 1.**
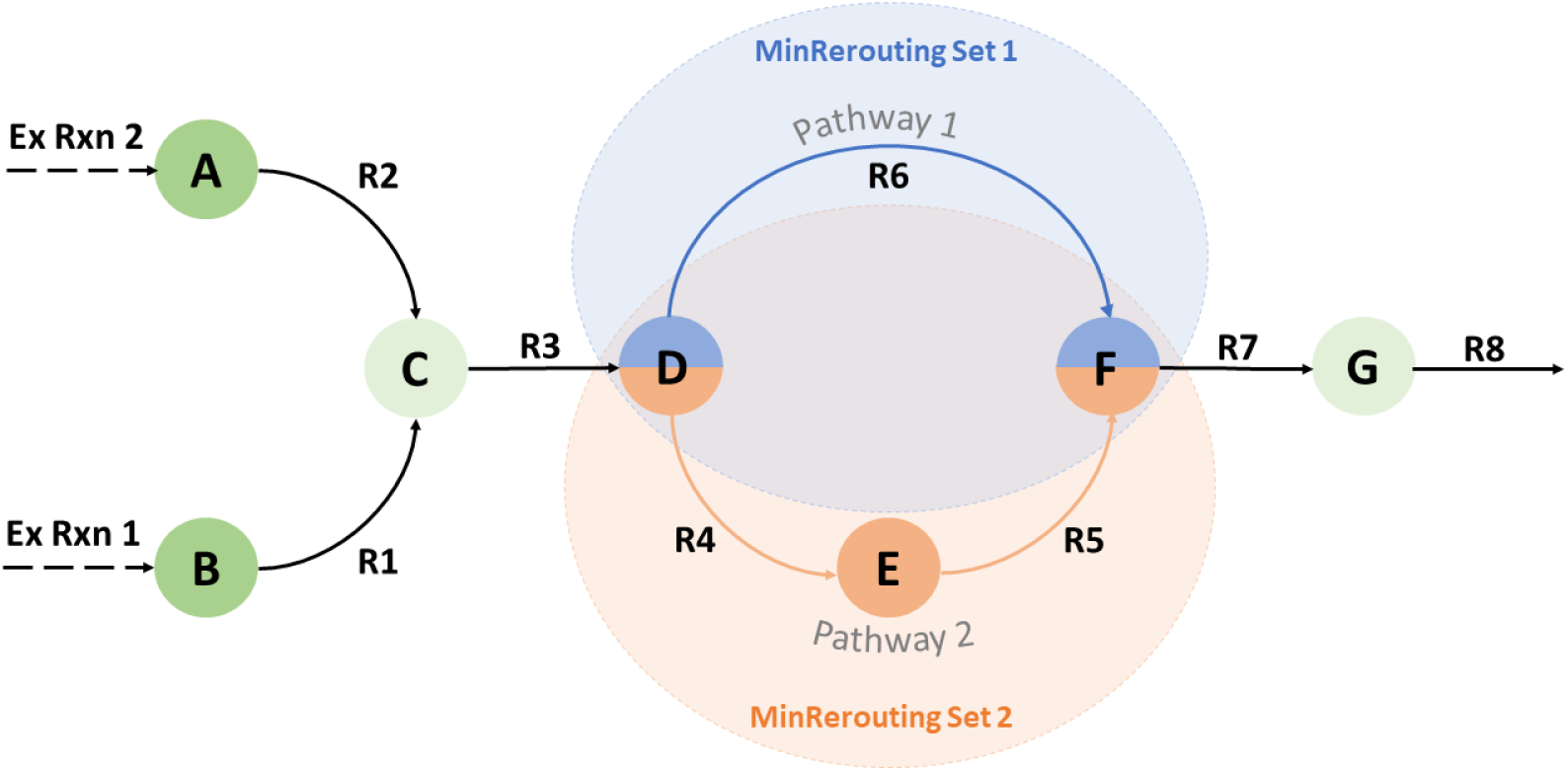
A sample metabolic network used to illustrate the concept of *minRerouting*. The nodes and edges are depicted as metabolites and reactions respectively. This metabolic network comprises 8 reactions, 7 metabolites, and 3 double lethal pairs. The *minRerouting* Sets 1 and 2 are highlighted using orange and blue rectangular boxes and the common reaction between the rerouting sets is *R*3.

The network consists of three double lethal pairs: {(*R*_1_*, R*_2_), (*R*_4_*, R*_6_) and (*R*_5_*, R*_6_)}. Taking the reaction pair (*R*_4_*, R*_6_) into consideration, we can see that when reaction *R*_4_ is active and *R*_6_ is deleted or inactive, all the fluxes will be routed through reactions *R*_4_ and *R*_5_. Similarly, when reaction *R*_6_ is active and *R*_4_ is deleted or inactive, all the fluxes will be routed through reaction *R*_6_. In addition to these changes, the fluxes routed through the remaining reactions could vary based on which pathway is chosen. For instance, the flux through reaction *R*_3_ could be significantly higher when the *R*_4_ pathway is used than when the *R*_6_ pathway is used.

Hence, when reaction *R*_4_ is active and reaction *R*_6_ is inactive or deleted, the rerouting set becomes *R*_3_*, R*_4_*, R*_5_. When reaction *R*_6_ is active and reaction *R*_4_ is inactive or deleted, the rerouting set becomes *R*_3_ and *R*_6_. The common rerouting set for the double lethal pair (*R*_4_*, R*_6_) consists of *R*_3_ and the complete rerouting set for the double lethal is (*R*_3_*, R*_4_*, R*_5_*, R*_6_).

In order to determine the minRerouting set, we first obtain the WT flux distribution, *v_W T_* , and the set of all lethal pairs in the model. Then for each lethal pair *R_i_* and *R_j_*, the optimal flux distribution *v*_Δ_*_Ri_* and *v*_Δ_*_Rj_* , that minimises the distance between the flux vectors is obtained, by an extension of the MOMA formulation.

The generalised *p*-norm formulation for obtaining the minRerouting of a model, for a given lethal pair, is as follows:

Step 1: An adaptation of MOMA is performed to obtain the optimal flux distributions *v*_Δ*Ri*_ and *v*_Δ*Rj*_ with minimal flux distance between them.

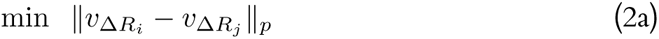

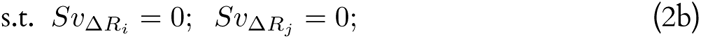

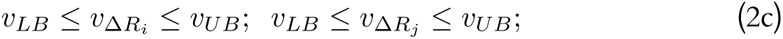

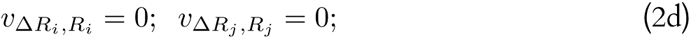

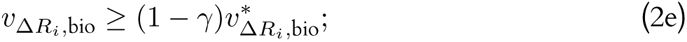

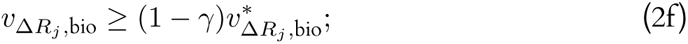

Step 2: The flux distributions *v*_Δ_*_Ri_* and *v*_Δ_*_Rj_* , obtained from Equation 2 are analysed. The reactions that have different flux values in *v*_Δ_*_Ri_* and *v*_Δ_*_Rj_* are identified as the rerouting set. The size of the rerouting set, the individual reaction flux difference and the total flux difference is also analysed.

Here, *v*_Δ_*_Ri_* and *v*_Δ_*_Rj_* represent the flux distribution when reactions *R_i_* and *R_j_* are deleted respectively. 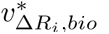 and 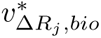 represent the optimal biomass flux in the models when reaction *R_i_* and *R_j_*are deleted respectively. *v*_Δ_*_Ri,Ri_* and *v*_Δ_*_Rj,Rj_* represent the flux through reactions *R_i_* and *R_j_* in the models where *R_i_* and *R_j_* are deleted respectively. *Γ* is the growth rate slack provided for the new flux distributions *v*_Δ_*_Ri_* and *v*_Δ_*_Rj_* from the optimal biomass 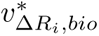 and 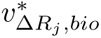

For obtaining the zero-norm solution, we use the LP formulation and IBM ILOG CPLEX v12.8 solver as it was one of the few solvers which supported zero-norm optimisation. The Gurobi solver is used for the one-norm and two-norm optimisation.

### 2.4 Parsimonious FBA (pFBA)

Parsimonious FBA (pFBA) [19] is a bi-level optimization problem, where first an optimal flux distribution that maximizes the biomass production is identified, followed by the minimization of total flux through all reactions. In addition to obtaining this flux distribution, pFBA also classifies all the reactions based on its enzymatic/metabolic efficiency. pFBA classifies the reactions into a total of six classes: Essential, ELE (Enzymatically Less Efficient), MLE (Metabolically Less Efficient), pFBA optimal, blocked, and zero flux reactions. These reaction classes are further used to classify the reactions in the lethal pairs and derive insights into the categorical distribution of these lethal pairs.

### 2.5 Flux Variability Analysis (FVA)

FVA [28] is used to obtain the maximum and minimum flux values that a reaction can carry in a model. FVA solves two LP problems (maximisation and minimisation) for each reaction in the model, while constraining the objective function (or) biomass growth rate value. FVA is formulated as follows:

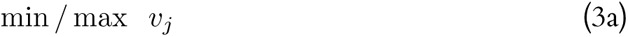

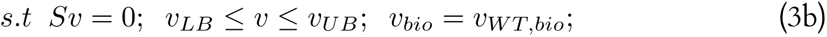

### 2.6 Plasticity and Redundancy in Synthetic Lethals

Previously [18], it has been suggested that synthetic lethal reaction pairs can be classified into two categories: Plastic Synthetic Lethals (PSL) and Redundant Synthetic Lethals (RSL). PSL comprises of reaction pairs where one reaction acts as a backup for the other, *i.e.* the second reaction becomes active when the first reaction is deleted. RSL comprises reaction pairs where both reactions are active simultaneously.

The classification approach proposed by a previous study[18] is based on flux vectors which are predicted using FBA. While an FBA solution satisfies all the flux and stoichiometric constraints for a given model, it only represents one possible flux instance from the permissible flux space. We propose a classification approach that is more systematic and thorough, taking into consideration the allowable flux space for each reaction that is part of a double lethal pair.

For instance, using the above approach, if the absolute fluxes of two reactions, reaction 1 and reaction 2, obtained from FBA, are greater than 0, then they are classified as RSL reactions. But, there is also a chance that reaction 2 can accommodate zero flux without any change in the optimal biomass flux, while reaction 1 is active. In this case, the reaction pair would have to be classified as a PSL pair. As the FBA only considers a single flux instance from the permissible space, we would not be able to correctly classify these reaction pairs. This necessitates a more thorough and systematic manner of classifying the reaction classes.

In order to classify the lethal pairs as PSL or RSL, we performed an FVA on the model. The product of the minimum and maximum flux ranges is used to determine the category of the reaction pair. Only when both the reactions are simultaneously active (with a positive or negative flux), while satisfying the biomass constraint, is the double lethal considered a RSL pair. The product of the sign of the minimum and maximum fluxes for RSL pairs includes the combinations [(*<* 0*, <* 0), (*<* 0*, <* 0)], [(*>* 0*, >* 0), (*>* 0*, >* 0)], [(*<* 0*, <* 0), (*>* 0*, >* 0)] and [(*>* 0*, >* 0), (*<* 0*, <* 0)]. In cases where this is not satisfied, a conditional FVA is performed before the double-lethal pair is classified as PSL or RSL.

For each of the ambiguous conditions, two FVAs are performed. In the first FVA, the flux of *R_i_* is constrained to be greater than 0 and in the second, the flux of *R_i_* is constrained to be less than 0. In this manner, the maximum and minimum fluxes of reaction *R_j_* are obtained when reaction *R_i_* is active. If the reaction *R_j_* can carry a flux value of zero, in either of the two constraint conditions, the reaction pair is considered to be a PSL reaction pair, as *R_j_* can be inactive when *R_i_* is active. However, if *R_j_* is always active under both the constraint conditions, the reaction pair is considered to be an RSL. We used this process to determine the classification of PSL and RSL classes instead of relying on a simple FVA because, in FVA, we obtain the maximum and minimum flux values of one reaction, independent of the activity of the other. The whole process is explained pictorially in Figure 2.

**Fig. 2.**
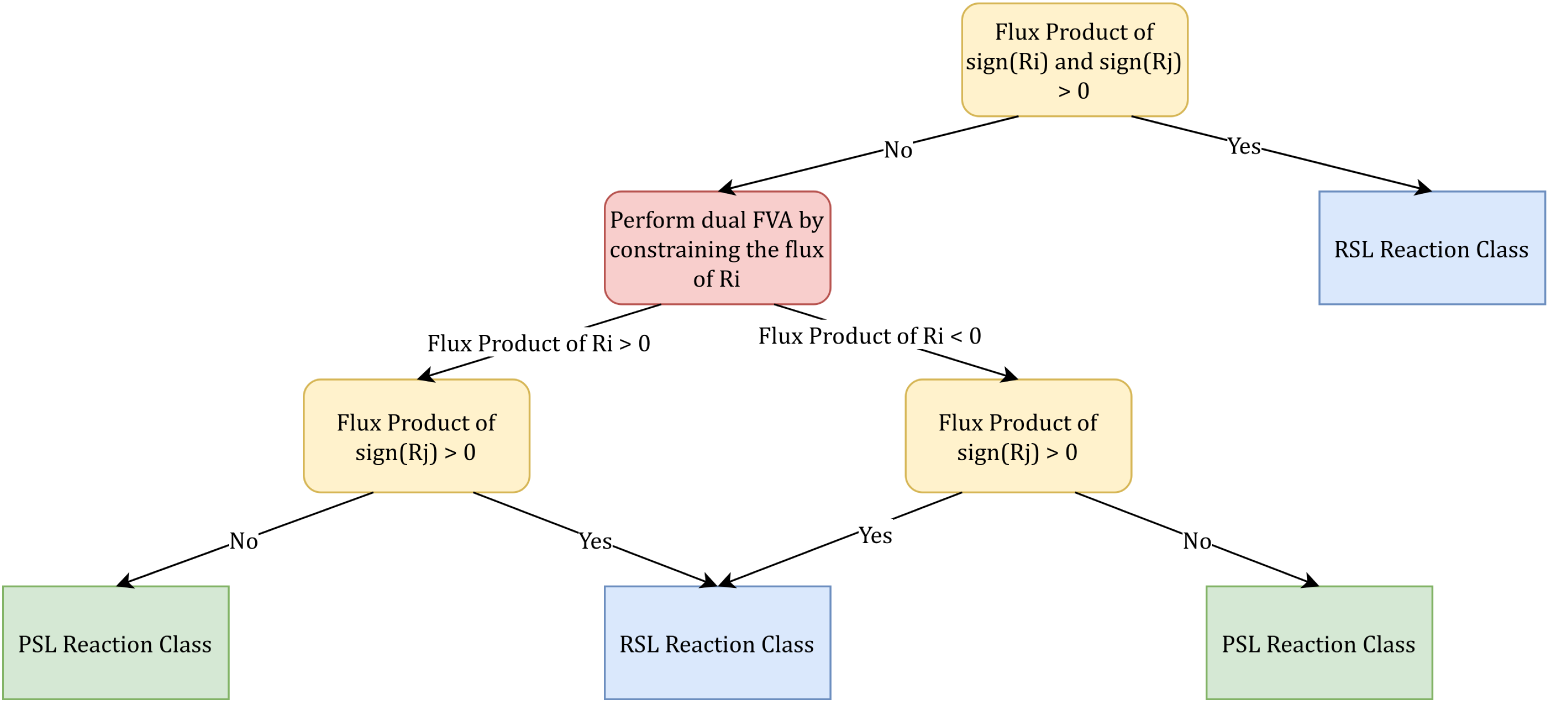
Flowchart depicting the classification of reaction pairs into PSL or RSL pairs. After the initial FVA, conditional FVAs are performed to classify the ambiguous reaction pairs. The reaction pairs are classified as RSL only when both reactions are simultaneously active.

#### 2.6.1 Number of Reactions Rerouted

For each synthetic lethal pair, the rerouting set would comprise two subsets — one for each of the double lethal reactions. The union of these two subsets would consist of the reactions with modified flux distribution. The reaction sets were analyzed, and the following properties were studied: (a) size of the SL cluster, (b) size of the common SL cluster, and (c) net difference between the flux vectors.

### 2.7 Metabolic Subsystem Analysis

A set of reactions that share a similar metabolic function is referred to as metabolic subsystem [29]. All the reactions in a Genome-Scale Metabolic Model are categorised under different metabolic subsystems. To identify the metabolic subsystem of the reactions that comprise a double lethal pair, the set of all distinct reactions to be analysed is obtained. Then, the subsystem of each of these reactions is obtained by systematically querying the BiGG database [30].

### 2.8 Implementation

The implementation of *minRerouting* and initial analysis were done using MATLAB. The metabolic cost analysis and flux rerouting analysis were done using Python and R. The figures were generated using R. The COBRA Toolbox [31] for MATLAB was used for all metabolic network analysis. All code written as a part of this project is open-sourced and can be accessed at https://github.com/RamanLab/minRerouting/.

## 3 Results

The analysis of the *minRerouting* sets has been carried out on eight genome-scale metabolic models. We have chosen organisms that represent key bacterial pathogens relevant to humans and are present in the BiGG database [30]. For *Escherichia coli*, found in the human gut, we have depicted two models, e_coli_core [32] and *i*ML1515 [33]. The model e_coli_core represents the simplified versions of only the most crucial pathways needed for its survival. The other models studied are for the bacteria *Helicobacter pylori*, *Klebsiella pneumoniae*, *Mycobacterium tuberculosis*, *Salmonella* Typhimurium, *Shigella sonnnei*, and *Yersinia pestis*. These models and the number of single and double lethal reactions predicted for them using Fast-SL are listed in Table 1.

**Table 1.**
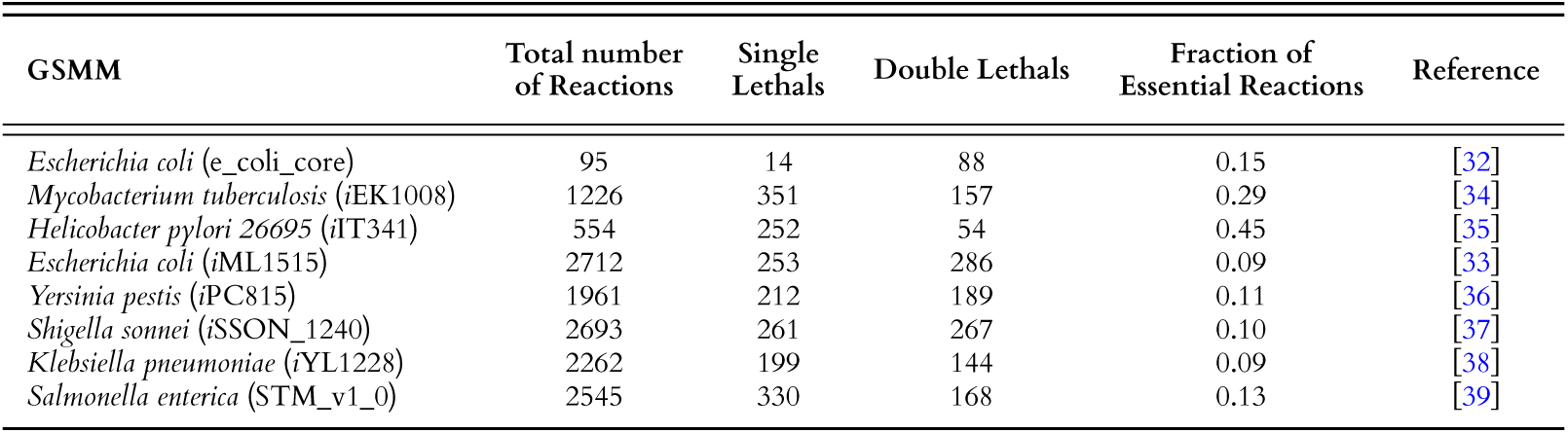
List of all the models analysed as a part of the study. The third and fourth columns indicate the number of single lethals and double lethal pairs identified using Fast-SL [26], [27]. The fifth column represents the fraction of reaction space composed of essential reactions.

### 3.1 Double lethals reveal metabolic redundancies

#### 3.1.1 Comparison of reaction submodules forming double lethals

Formerly, two studies [17, 18] found the presence of double lethals in the species *E. coli*, *M. pneumoniae*, *S.* Typhimurium, and *S. sonnei* using FBA. Notably, our work is consistent with their results regarding the number of synthetic double lethals obtained for different species and their composition of reactions. For the *E. coli* model *i*JO1366, 254 synthetic lethals [18], and 229 synthetic lethals [17] were found, while we found 286 synthetic lethals in the model *i*ML1515.

From Table 1, we see that the distribution of the number of synthetic single and double synthetic lethals is distinct for the two species, *M. tuberculosis* and *H. pylori* , which are known to be specially adapted to their host environments compared to the other species [40, 41]. They exhibit limited metabolic diversity, depending primarily on a single method of energy generation. Reactions associated with this method are indispensable to these organisms. Thus, their genome-scale metabolic network encompasses 29% and 45% of essential reactions. In contrast, generalists with multiple energy production pathways do not focus on one pathway as critical or essential to their survival and display a lower proportion of reactions essential to their networks. This parallels previous observations [4, 18] that specialists have more essential genes, which are robust against perturbations mainly because of gene duplications and not alternate pathways. Our results also show that the double lethals for the two specialists are less than 50% of the single lethals present in the reaction space, implying a lesser reliance on redundancies for robustness.

Figure 3 shows that more than 500 double-lethal pairs are present in at least one organism, indicating the high level of redundancy present across metabolic networks. Among these, few are shared across at least five of the models analysed in the study. The tendency of a reaction pair to appear in multiple species with different adaptations implies that these could be potential super targets for drug therapies. The common synthetic lethal pairs examined across the organisms are all from the Pentose Phosphate Pathway. The second most common reactions are from the glycolytic pathway and different amino acid biosynthesis pathways. Similar to observations made by Barve *et al*.[42], the reactions that were essential belonged to linear or anabolic pathways such as ATP and histidine synthesis while redundancies were present only in more reticulate pathways such as the pyruvate or glycoysis metabolic pathways.

**Fig. 3.**
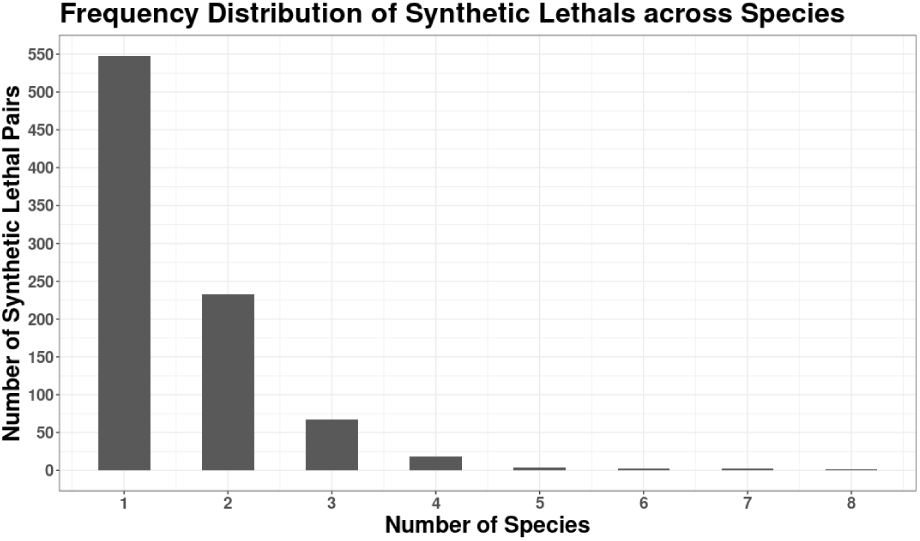
Distribution of common double lethals across models. 500 reaction pairs are unique to the model while very few pairs are present across all the eight models.

A broader understanding is obtained by looking at each organism’s submodule distribution of double lethals, as shown in Figure 4 and Figure 5 for the models *i*ML1515 and *i*EK1008. While it is intuitive to think the reaction pairs would arise from the same submodule, for all models, we see that at least 50% of the reactions are from different sub-modules, *i.e.*, the synthetic lethals are inter-pathway. This could be because of the ripple effect caused by deleting one reaction, which causes small changes in all other connected pathways. Inter-pathway synthetic lethals also highlight an organism’s need for cross-talk between pathways, such as energy production and nucleotide metabolism [43].

**Fig. 4.**
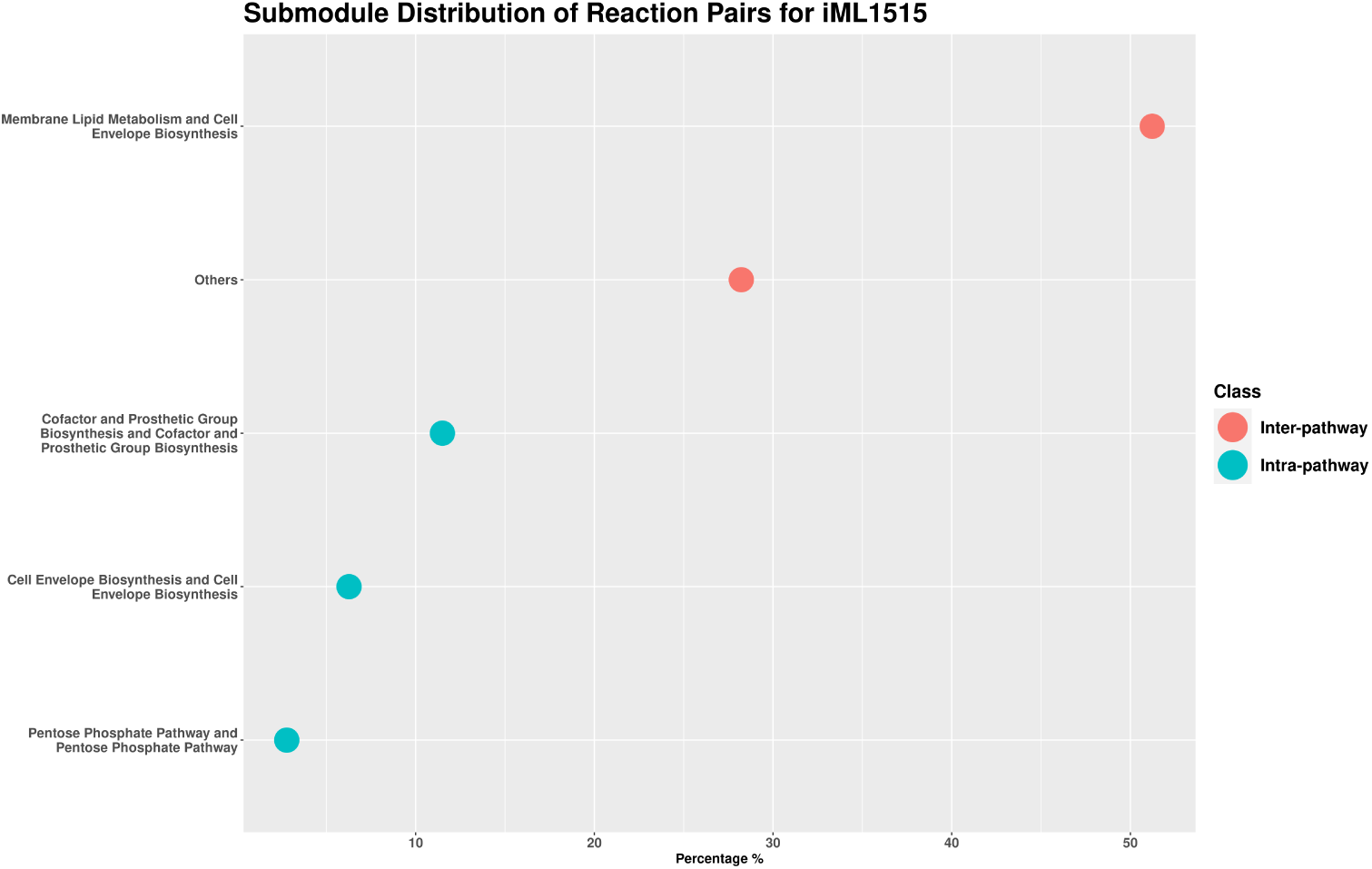
Metabolic Subsystem Analysis for the model *i*ML1515. The distribution shows that more than half the lethal pairs are from differing submodules. Only the submodules occurring more than the mean of the distribution are depicted for clarity.

**Fig. 5.**
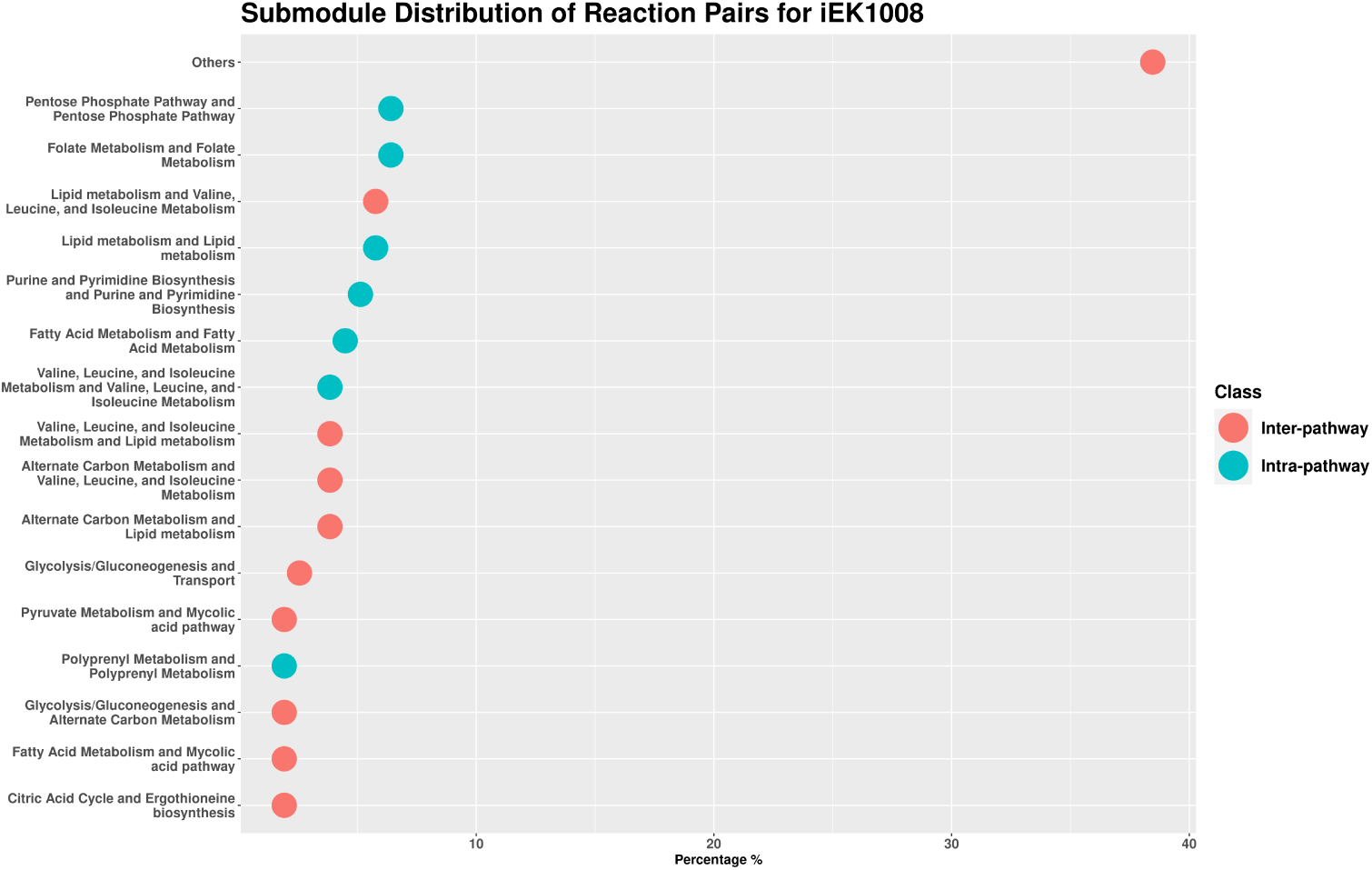
Metabolic Subsystem Analysis for the model iEK1008. The distribution shows lethal pairs from unique submodules such as mycolic acid pathway and pyruvate metabolism. Unlike *i*ML1515, Cell Envelope Biosynthesis does not appear in any of the top pairs. Only the submodules occurring more than the mean of the distribution are depicted for clarity.

In simpler and closely related bacterial systems, such as *E. coli* and *S. sonnei* , more than 50% of synthetic lethal reactions involve cell envelope biosynthesis, where membrane lipid metabolism reactions act as their backups. These systems also exhibit reaction pairs from the cofactor and prosthetic group biosynthesis submodule. *Y. pestis*, *K. pneumoniae*, and *S.* Typhimurium, phylogenetically different from the above two species [44], additionally had reaction pairs from submodules associated with amino acid metabolism and glycerophospholipid metabolism. Finally, the specialist species *M. tuberculosis* and *H. pylori* have distinct dominant submodules, such as mycolic acid production and heme transport, respectively. The distributions for the other six models are given in the Supplementary Results presented in Appendix A. Thus, the redundancies are organism-specific, with double lethal reactions from submodules required for biomass synthesis, directly or indirectly. Some submodules, like the Pentose Phosphate Pathway, are shared in all organisms. These findings align with those from earlier experiments [10, 17, 18].

#### 3.1.2 Tendency of reactions in forming double lethals

We next define the Redundancy Index (RI) as the number of times a reaction is repeated in synthetic lethal pairs within a species. Several studies indicate that pathways lacking redundancy tend to have more critical functions for survival than those with redundant pathways [43, 45]. However, there is evidence contradicting this notion [10, 46]. Specifically, when a reaction is engaged in multiple pairs as a synthetic lethal and consequently possesses multiple backups, its function likely plays a significant role in ensuring the organism’s survival. From Figure 6, we observed that the models had a mean RI in similar ranges despite having different numbers of reactions and different adaptations. The reactions with a high RI were from different submodules for each organism, ranging from central carbon metabolism to transport and, more specifically, lipid metabolism for the bacterium *M. tuberculosis* . As defined beforehand [42], a reaction is essential if its elimination turns off biomass production, which is indicated by the production of fatty acids, amino acids, and purine/ pyrimidine. Thus, these submodules occurred more often in the single lethals list, and reactions that had redundancy in the form of double lethals were albeit important but impacted biomass production indirectly. The *minRerouting* approach reveals an essential characteristic of metabolic networks, i.e., the inter-dependencies of the pathways to produce biomass are revealed. These inter-dependencies are due to molecules with a high RI acting as connecting points for other pathways in a network. Suthers *et al*. [43] investigated the topological structure of redundancy and observed comparable findings regarding the essentiality of various reaction modules.

**Fig. 6.**
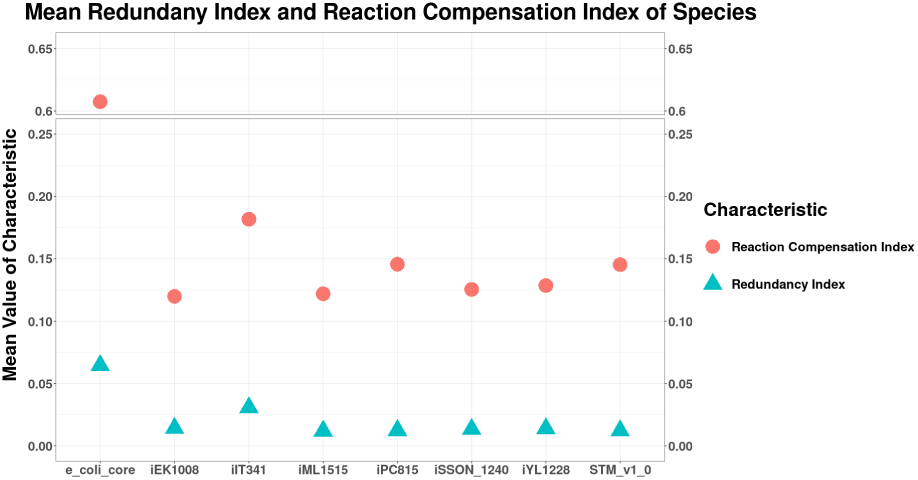
Mean Redundancy Index and Reaction Compensation Index of Reactions forming Synthetic Lethal pairs, across Organisms. Reactions from e_coli_core have a higher RCI and RI compared to other models.

The model e_coli_core exhibits a notably higher RI, approximately ten times greater than other models, owing to its inclusion of only the essential reaction pathways. The smaller number of reactions in it leads to a high level of redundancy since it necessitates that the few genes become backups for each other. Specifically, the model has 88 double lethals among only 95 reactions, implying extensive backing of the core metabolic pathways. These pathways include central carbon metabolism, cell membrane synthesis, transport, and amino acid biosynthesis. Varying the levels of complexity of a model results in a change in which reactions have backups.

Similarly, the propensity of a reaction to be part of a synthetic cluster, or its Reaction Compensation Index (RCI), has been tested and shows similar trends as RI in terms of value, seen in Figure 6. However, the reactions that have a high RI do not necessarily have a high RCI. A high RCI indicates reactions that are important for all the key pathways that the synthetic clusters form but these are not necessarily essential at least at the order of a double lethal.

To further understand how these reactions compensate for each other by rerouting flux, and ensure their viability, we further analysed the results from the *minRerouting* algorithm.

### 3.2 Flux Redistribution Analysis

For each of the *p−*norms, the resultant *minRerouting* set is analysed. The characteristics of the *minRerouting* set, which are studied are as follows:

#### 3.2.1 Size of the *minRerouting* Set

The size of the minimal rerouting set, as discussed in the Introduction, is the number of reactions through which flux is rerouted. *minRerouting* allows the user to input their preferred norm to minimise the rerouted flux since differences in the calculation of the norms result in slightly different results. *l*_2_ norm is called the least squares error norm as it minimises the sum of the square of the differences. As a result, it tends to be influenced by outliers which can lead to unexpected solutions. However, its solution is unique and stable. On the other hand, the *l*_1_ norm gives a sparse solution, but with the possibility of multiple solutions for the minimisation problem. The *l*_0_ norm, computationally difficult to compute also results in a sparse solution. The comparison of the *minRerouting* cluster size across three norms, shown in Figure A4 thus reveals that the *l*_0_ norm optimisation results in the smallest SL Cluster Size, followed by the *l*_1_ and *l*_2_ norms, respectively.

Interestingly, the sizes of the clusters in *l*_1_ norm are lesser than observations made by Massucci *et al*. [17] as seen in Figure A12. In stressful environments, restructuring metabolism incurs functional and structural costs. Our approach minimises the alterations in flux required, even if it means a partial reduction in growth rate, such that the expenses are reduced. Thus, a smaller cluster size is expected. The cluster size for each double lethal pair of an organism is shown in Figure 7 along with other properties of the cluster for easy comparison across models.

**Fig. 7.**
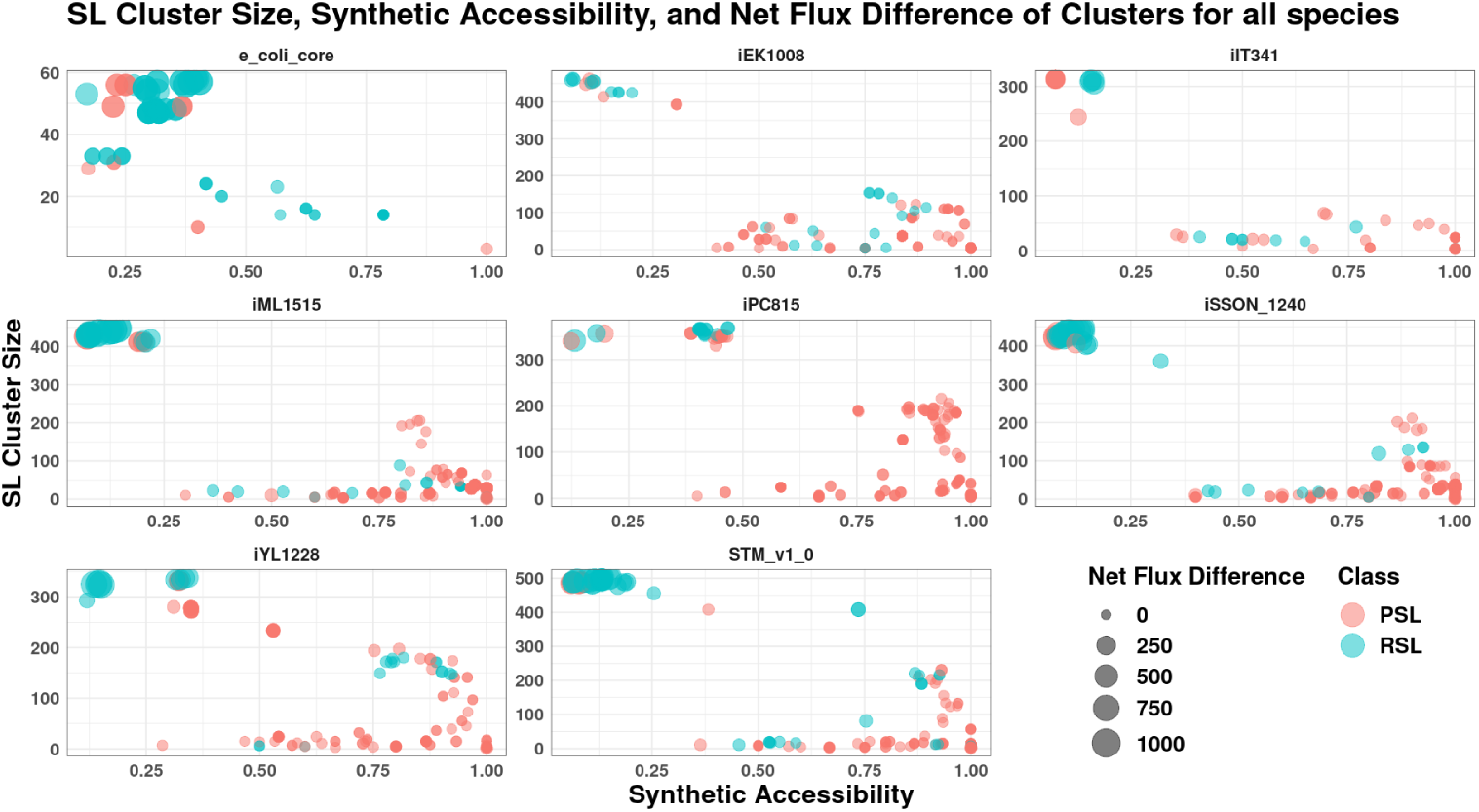
Cluster Size, Synthetic Accessibility, and the Net Flux Difference between the two metabolic states of Reaction Pairs. Mostly, the pairs that have high synthetic accessibility have a low net flux difference and small cluster size. Outliers showing different properties exist in the top left corner for all the models except e_coli_core.

#### 3.2.2 Size of the Common *minRerouting* Cluster Set

The size of the common minimal rerouting set, as discussed in the Introduction, is the number of common reactions through which flux is rerouted. The comparison of the common *minRerouting* Set size across norms is shown in Figure A4.

Once again, the results obtained show that the *l*_0_ norm optimisation results in the smallest Common SL Cluster Size, followed by the *l*_1_ and *l*_2_ norms, respectively. In certain models, the median Common SL Cluster Size is 0. This indicates that the *l*_0_ norm, in addition to ensuring minimum SL Cluster size, forces the two flux vectors to take completely exclusive reaction pathways, with no reaction (with modified flux) overlap. Since the organism undergoes complete rewiring, the number of new reactions flux is rerouted through is termed Synthetic Accessibility. The organism has to divert its energy into activating/ deactivating the reactions not common between the two reaction pair deletions. A Synthetic Accessibility of 1 would imply no common reactions between the metabolic states of a synthetic lethal pair. While the range of Synthetic Accessibility is vast, on average, at least 25% of reactions in a cluster are turned on/off when switching states. A low Synthetic Accessibility could mean that the function of the reaction is not being completely replaced but is being compensated for by a collective group of reactions mediated through its pair. A high Synthetic Accessibility could mean that most of the reactions that get activated in the mutant are not needed in the wild-type state. In literature, there are multiple schools of thoughts for the role of redundancy in metabolic networks. True redundancy is believed to not be possible as it is evolutionarily unstable [47, 48]. A genuinely redundant gene coding for the redundant reaction would be lost completely through genetic drift since its mutation would incur no fitness costs to the organism. Hence, we see that synthetic accessibility indeed has a vast range. However, the clusters with a small cluster size tend to have a high Synthetic Accessibility. Thus, the backing is efficient and reactions are not unnecessarily active. However, one group of clusters is unique as seen in Figure 7. They are clusters that have a high cluster size and a low Synthetic Accessibility as seen in the top left corner of every model. Thus, Synthetic Accessibility gives us an idea of the efficiency of the metabolic network and the cost of replacing that function that the organism is willing to bear, even with a decrease in its growth rate.

#### 3.2.3 Net Flux Difference

The flux difference is the total flux difference between the two flux vectors representing the deletion of individual reactions that make up the double lethal. The comparison of the net flux difference across norms is shown in Figure A5. Flux difference shows the same trend for *l*_0_, *l*_1_, and *l*_2_ norms. When comparing with Massucci *et al*. [17], the net flux difference between reaction pairs is significantly lesser when comparing our *l*_1_ norm results with theirs for the organism *E. coli*, *S. sonnei* , and *S.* Typhimurium as seen in Figure A12. The flux difference gives an indication of the cost of maintaining the redundancy between the reaction pairs. *minRerouting* thus reveals the flux redistribution that the organisms can undertake to decrease this cost further. Since the net flux difference is comparable between the norms while the cluster size is more for *l*_2_, the average change in a reaction flux is lower for *l*_2_ norm than *l*_1_ and *l*_0_ norm. This may be because the *l*_2_ norm penalises differences more as it squares them to obtain the error.

The average flux difference of clusters is high in the e_coli_core model, due to its limited ability to adjust to perturbations. Thus, even though organisms do not need many reactions to exist, more reactions provide robustness to the network.

The outlier group for all models barring e_coli_core, mentioned previously, consisting of clusters with a low Synthetic Accessibility and high cluster size also have a high net flux difference which can be seen in Figure 8. From the bubble plot, we see that the outliers exist as two subgroups in each species. The outliers that still have a lesser flux difference all have higher Synthetic Accessibility and lesser cluster size. The ones with higher are the second group with lower Synthetic Accessibility and bigger cluster size. The Pentose Phosphate Pathway module and reactions from various transport submodules show up in many of the organisms along with species-specific reactions. In *K. pneumoniae*, we see histidine metabolism as a replacement for the TCA cycle as a major subgroup of the outliers. For *M. tuberculosis* we see the reaction space from mycolic acid metabolism and for *H. pylori* , reactions from the urea metabolism are present in the outliers. These outliers are important reactions of interest since they are not bypassed even though their deletion causes a major disruption in the flux distribution of the organism.

**Fig. 8.**
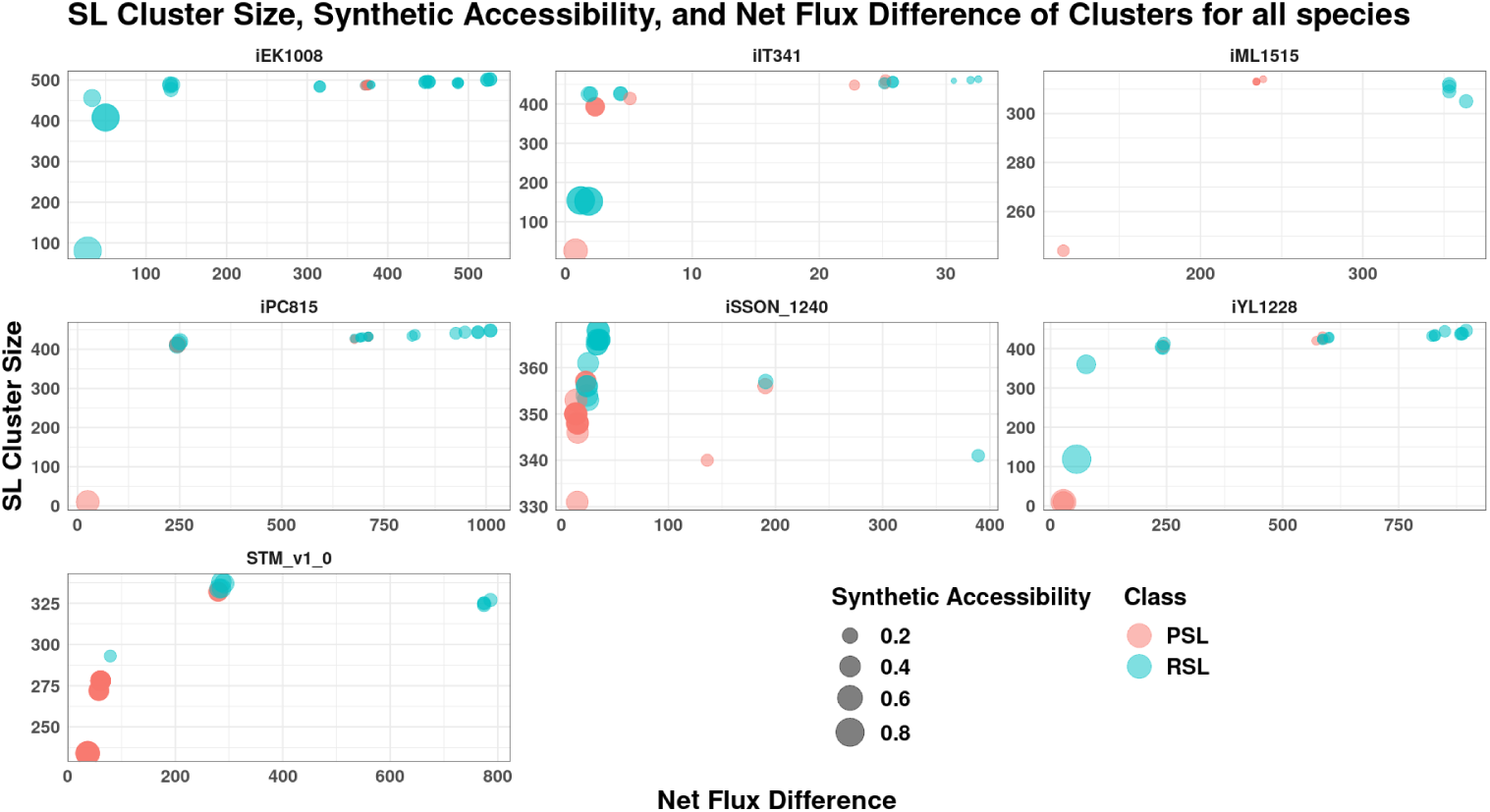
Cluster Size, Synthetic Accessibility and the Net Flux Difference between the two metabolic states of Outliers across species. Here, we can see that RSLs have the highest flux difference owing to their cluster size. Pairs with a higher synthetic accessibility have lesser flux differences despite large cluster sizes.

The rerouting also occurs with two strategies-PSLs and RSLs. What difference does the plasticity or redundancy make to the observations we have seen? We next check how they impact the robustness of metabolism.

### 3.3 Metabolic Efficiency Analysis

#### 3.3.1 Overall Distribution

Using Flux Variability Analysis and the methodology described above, we have classified the clusters into Redundant Synthetic Lethals (RSLs) and Plastic Synthetic Lethals (PSLs). This classification of synthetic pairs is based on the two strategies taken by the synthetic pairs to maintain robustness. It helps explain different properties seen till now concerning cluster size, flux difference, and composition. Previous analysis [17, 18] has revealed that PSLs are the more complicated way of acquiring redundancy, needing sophisticated functional organisation with fewer resources, while RSLs represent a more rudimentary strategy. For simple species like *E. coli*, their results showed that RSLs had more intra-pathway submodule pairs, PSLs had more inter-pathway submodule pairs, and for *M. pneumoniae*, no pattern was observed for RSLs or PSLs. From Figure 9, we see that the number of RSLs is higher than PSLs only in the e_coli_core model, which is the simplified metabolic network of *E. coli* without any functional or structural complexities of metabolic networks. Thus, the organisms tend to promote the plasticity of networks. The clusters do not prefer inter-pathway or intra-pathway reaction pairs but seem to depend more on the organism and which pathways are important to it. For organisms like *E. coli*, the most common submodules for pairs were inter-pathway due to their dependency on the cell envelope synthesis pathway and the membrane lipid biosynthesis pathway.

**Fig. 9.**
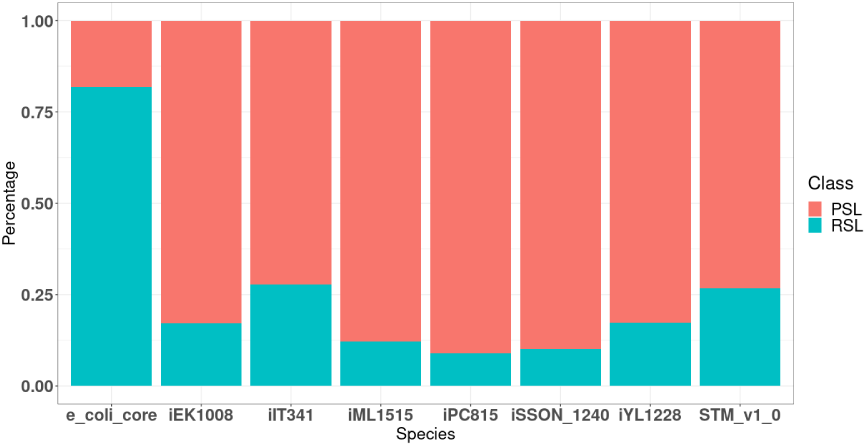
PSL-RSL distribution across models. While in most models the fraction of PSLs are much greater than that of RSLs, the trend is reversed in the case of the model e_coli_core. This could be expected because e_coli_core only comprises the metabolic core of *E. coli*, and hence, most of the double lethal pairs comprise reactions that require to be simultaneously active.

From Figure A13, PSLs exhibit a smaller cluster size for each species than RSLs, as depicted in the figure. This smaller cluster size indicates that transitioning between two states is structurally simpler, with fewer changes needed. This ease of transition is also reflected in the functional costs, as PSLs have a lower flux difference between the two states as seen in Figure A15. The difference in costs and efficiency is highlighted by the higher Synthetic Accessibility of PSLs as compared to RSLs in Figure A14. These trends are again not seen in e_coli_core, which is unique due to limited flexibility stemming from its reactions and connectivity.

While the RSLs and PSLs help us understand the above trends, why does the network have differing strategies? Why don’t the RSL reaction pair compete against each other, with only one reaction becoming evolutionarily stable? In PSLs, why does the inactive reaction stay in the network when it is not needed in wild-type scenarios? To understand why this happens and dive deeper into the types of reactions that contribute to RSL and PSL pairs, the metabolic efficiency analysis of the reactions was performed using the reaction classes returned by pFBA [19].

#### 3.3.2 Model wise distribution

Firstly, we see how the type of reaction, pFBA Optima, blocked, enzymatically less efficient (ELE), metabolically less efficient (MLE), zero flux reaction, or essential reactions, influenced double lethal pair formation. From Figure A2, we see that the Redundancy Index does not depend on the type of reaction. In fact, we see a high composition of ELE and MLE reactions to be redundant which is unexpected yet concurrent with the study by Wang *et al*. [46]. We also see that the percentage composition of RSL pairs is lesser than PSLs, possibly because of the more evolved nature of PSLs. Yet, the RSLs are found to be composed of mainly pFBA Optima reaction pairs (Figure 10, Figure 11). In PSLs, while one of the reactions was pFBA Optima (possibly the active reaction in wild type state), the other reaction was seen to be part of any of the other types of reactions. The reason reactions other than pFBA Optima form redundant pairs in nature is puzzling, but can be explained by various evolutionary forces acting simultaneously over the organism while under the influence of the environment [46].

**Fig. 10.**
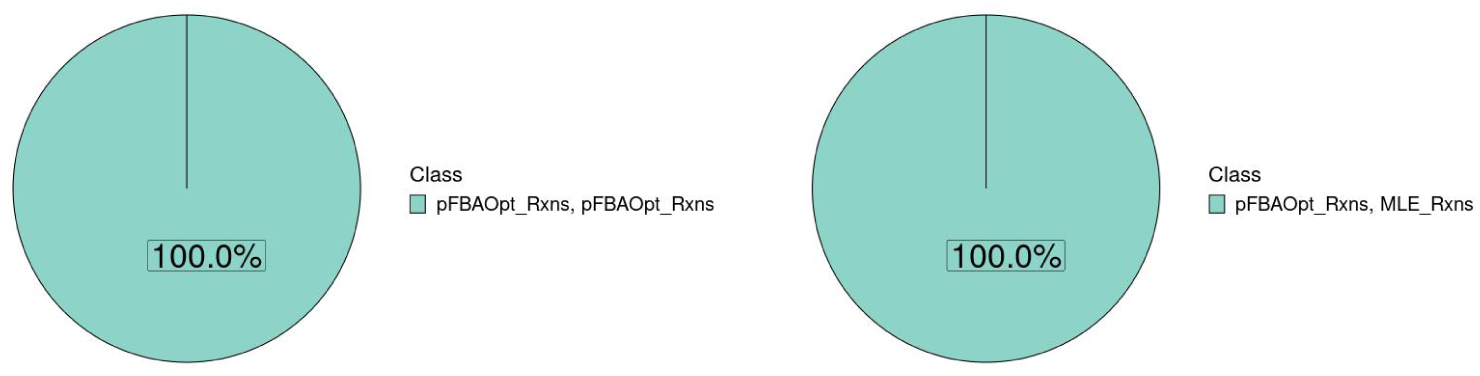
Schematic of reaction pair distribution between the RSL and PSL classes for e_coli_core. Majority of the reaction pairs that are classified as RSL pairs are (pFBA optimal, pFBA optimal) pairs as seen in the left sub figure. The distribution for the rest of the models can be accessed in the supplementary results provided in Appendix A

**Fig. 11.**
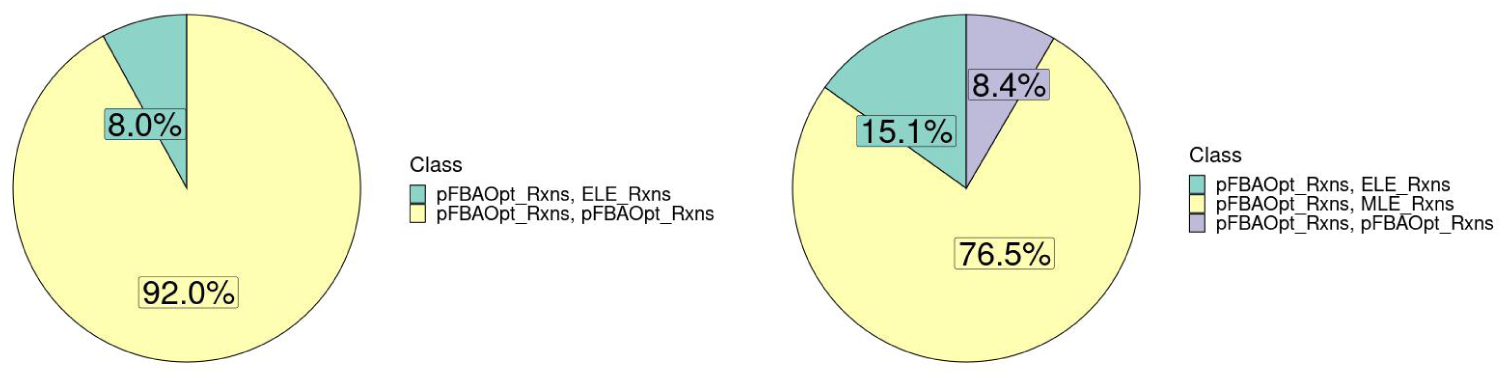
Schematic of reaction pair distribution between the RSL and PSL classes for iYL1228. Majority of the reaction pairs that are classified as RSL pairs are (pFBA optimal, pFBA optimal) pairs as seen in the left sub figure. The distribution for the rest of the models can be accessed in the supplementary results provided in Appendix A.

From the metabolic efficiency analysis, it is clear that the type of reaction influences the formation of different classes of synthetic lethals and in turn, its properties. Comparisons between e_coli_core and *i*ML1515 also reveal that the extent of connections and density of the network also impact the rerouting of metabolism. These results indicate how the robustness of metabolic networks is more complex than anticipated.

## 4 Discussion

In this study, we have looked at how organisms, especially pathogenic bacteria, rely on the presence of redundancy in the form of synthetic lethals to make themselves robust against genetic perturbations. With the help of synthetic lethals and our proposed algorithm, *minRerouting*, we have analysed how flux is redirected in metabolic networks upon perturbations. While we see that the specialist bacteria studied here, *M. tuberculosis* and *H. pylori* do not prefer the use of alternate pathways as a strategy for robustness, other species rely on double synthetic lethals for various kinds of functions. Yet, there is a set of synthetic lethals that are common across species. These are from the Pentose Phosphate Pathway. Reactions that are from these common pathways are important as their functionalities could be potential targets for therapy and antimicrobial resistance. They represent nodes in the network that interact with multiple other nodes simultaneously.

Our optimisation formulation, *minRerouting*, is used to obtain the minimal set of reactions through which fluxes are rerouted. We used three different approaches based on the norms - sparse (*l*_0_), linear (*l*_1_) and quadratic (*l*_2_) approaches and obtained different *minRerouting* sets for each of the norms. It is likely that the *l*_0_ captures the final steady-state of the cell post-adaptation to the knock-out, while the other norms could capture the transient response of the cell to the perturbation, *i.e.* gene deletion, similar to those observed by Shlomi *et al*. [49].

We showed that the size of the *minRerouting* set is the smallest when the sparse formulation is used and learnt that sparsity also forces the use of exclusive reaction rerouting, which results in a null common rerouted reaction set Figure A4.

We also proposed a systematic, conditional FVA approach to classify double lethal pairs into PSL (Back-Up Reactions) or RSL (Parallel-Use Reactions) reaction pairs. Parallel to remarks made previously[18], the Synthetic Accessibility of PSLs highlights their sophisticated nature in contrast to RSLs and in all models barring the e_coli_core, the PSLs were higher in number. These RSL and PSL pairs have differing properties in terms of the number of reactions through which flux is rerouted, the new set of reactions that become active when flux is rerouted, as well as the costs of rerouting. We explored the reasons for such a disparity and hypothesised that the RSL Pairs should have higher metabolic efficiency as both the reactions are simultaneously active, while reactions in the PSL Pairs would have lower metabolic efficiency. The results of our study have proved that this was indeed true. The RSL Reaction Pairs of most of the organisms (excluding *Shigella sonnei*), comprise completely (or majorly) of (*pFBAOptimal, pFBAOptimal*) reaction pairs. As *pFBAOptimal* reactions are considered crucial for the growth of an organism, our hypothesis was validated. Reactions comprising PSLs must have come about by various evolutionary processes such as horizontal gene transfer, and pleiotropy and may not have been a result of back-up for adaptation or robustness. These observations were made possible by the results from the *minRerouting* algorithm.

An extension of the *minRerouting* can be used to understand the complex metabolic reroutings that occur in several diseases. Particularly, in the case of cancer, where the cells re-programme their metabolic activities, rerouting fluxes in such a way that they can continue to proliferate and maintain their malignant properties, *minRerouting* can help us understand these reroutings and perhaps help in finding better therapeutic cures. Potential synergistic effects of drugs can be unearthed from applications of this algorithm. The submodule distribution and the metabolic efficiency results take us one step closer to understanding the structure of redundancy present across metabolic networks. They reveal hidden dependencies between reactions and the influence they have on flux rerouting in a network. This approach of interpreting flux switching is crucial to understanding redundancies in metabolic networks and their differing roles.

**Supplementary information.** Supplementary information has been provided in Appendix A.

## Declarations

- Funding TM thanks IBSE for the post-baccalaureate fellowship. The other authors received no specific funding for this work.
- Conflict of interest/Competing interests The authors declare no financial or non-financial competing interests.
- Ethics approval and consent to participate Not Applicable
- Consent for publication Not Applicable
- Data availability All models analysed during the study are taken from the BiGG database [30].
- Materials availability Not Applicable
- Code availability The ‘minRerouting’ algorithm is available from https://github.com/RamanLab/minRerouting
- Author contribution KR conceptualised the study. JSN and KR designed the investigation. TM, OKM and SM ran the computational experiments. TM, OKM, SM and KR interpreted the results. KR supervised the project. SM, OKM and TM drafted the initial draft of the manuscript. All authors read and approved the final manuscript.

## Appendix A Extended Data

The reaction distribution is explained pictorially in Figure A1.

**Fig. A1.**
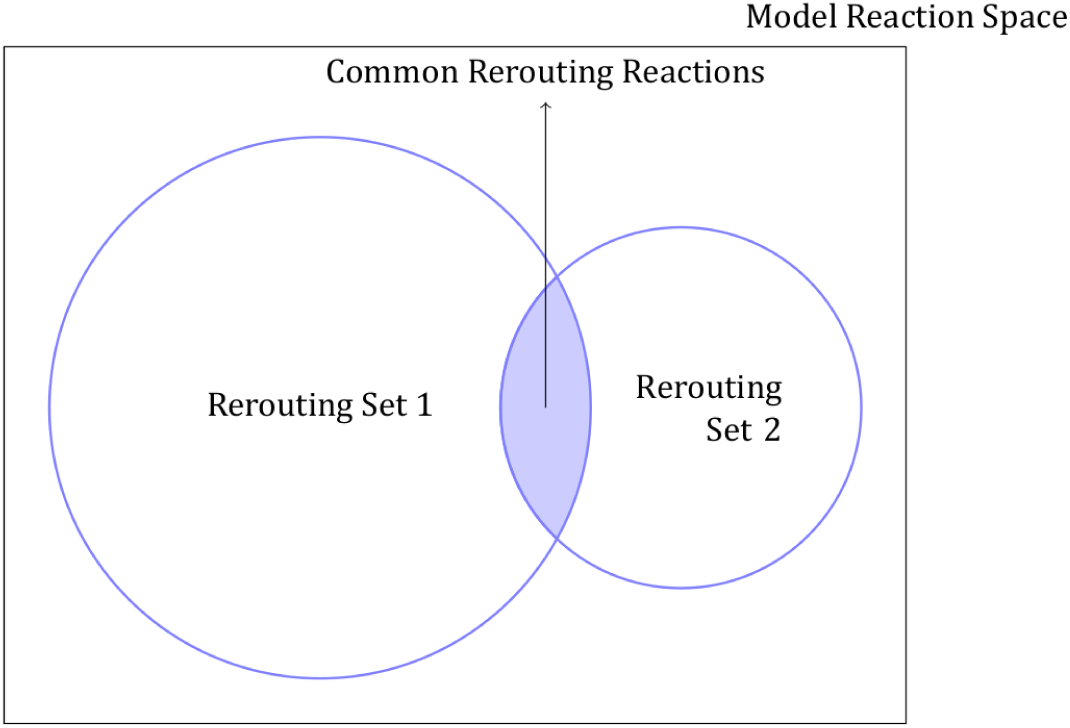
Representation of the Reaction Space and the different Rerouting Sets.

**Fig. A2.**
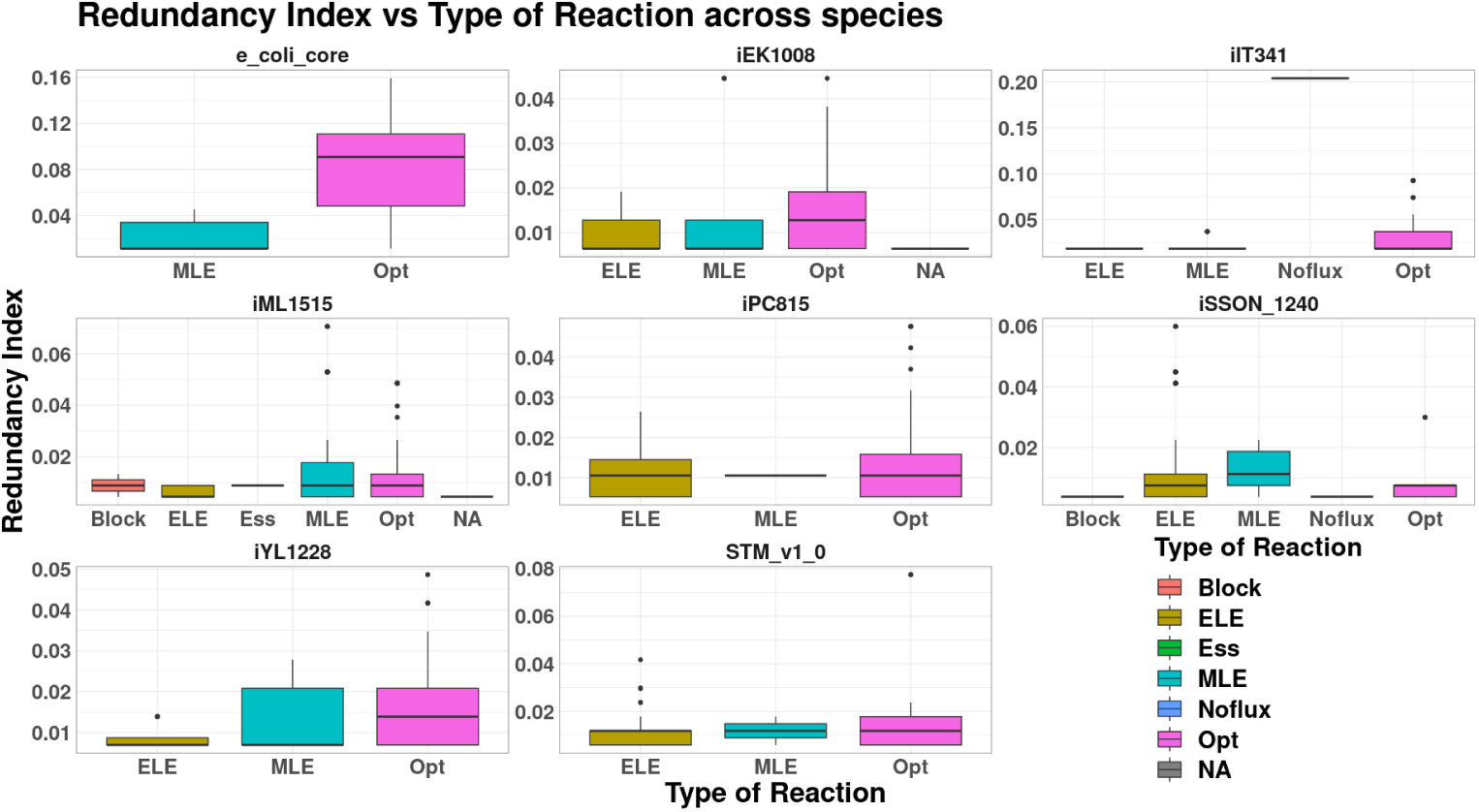
Redundancy Index of Reactions distributed over the type of reaction for each species. There is no relation between the type of reaction and its tendency to form redundant pairs.

**Fig. A3.**
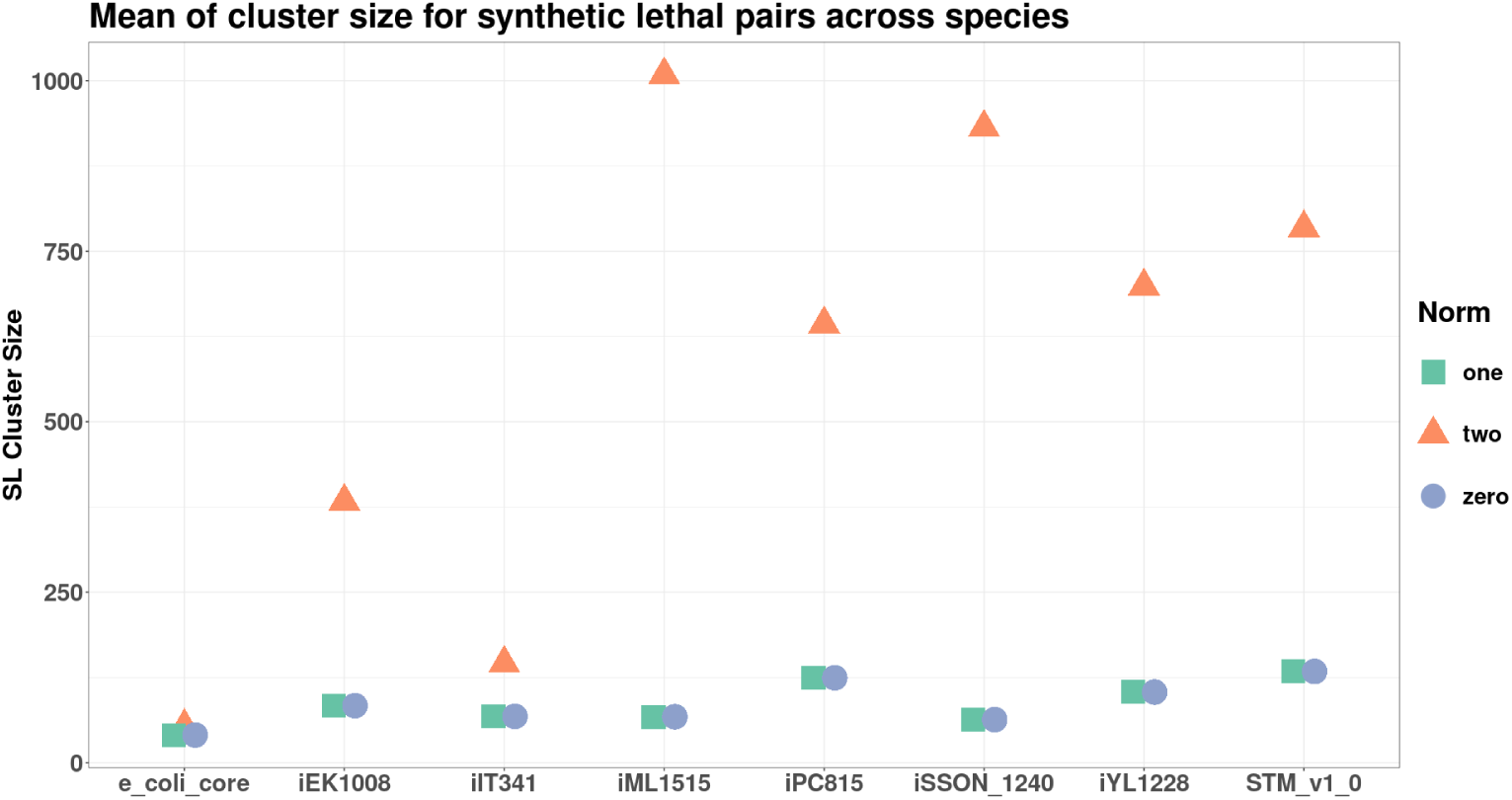
Distribution of SL Cluster Size across Organisms and Norms. Notice that the SL Cluster Size is the smallest for the zero and one norms, while the SL Cluster Size obtained using two norm is approximately an order of magnitude higher.

**Fig. A4.**
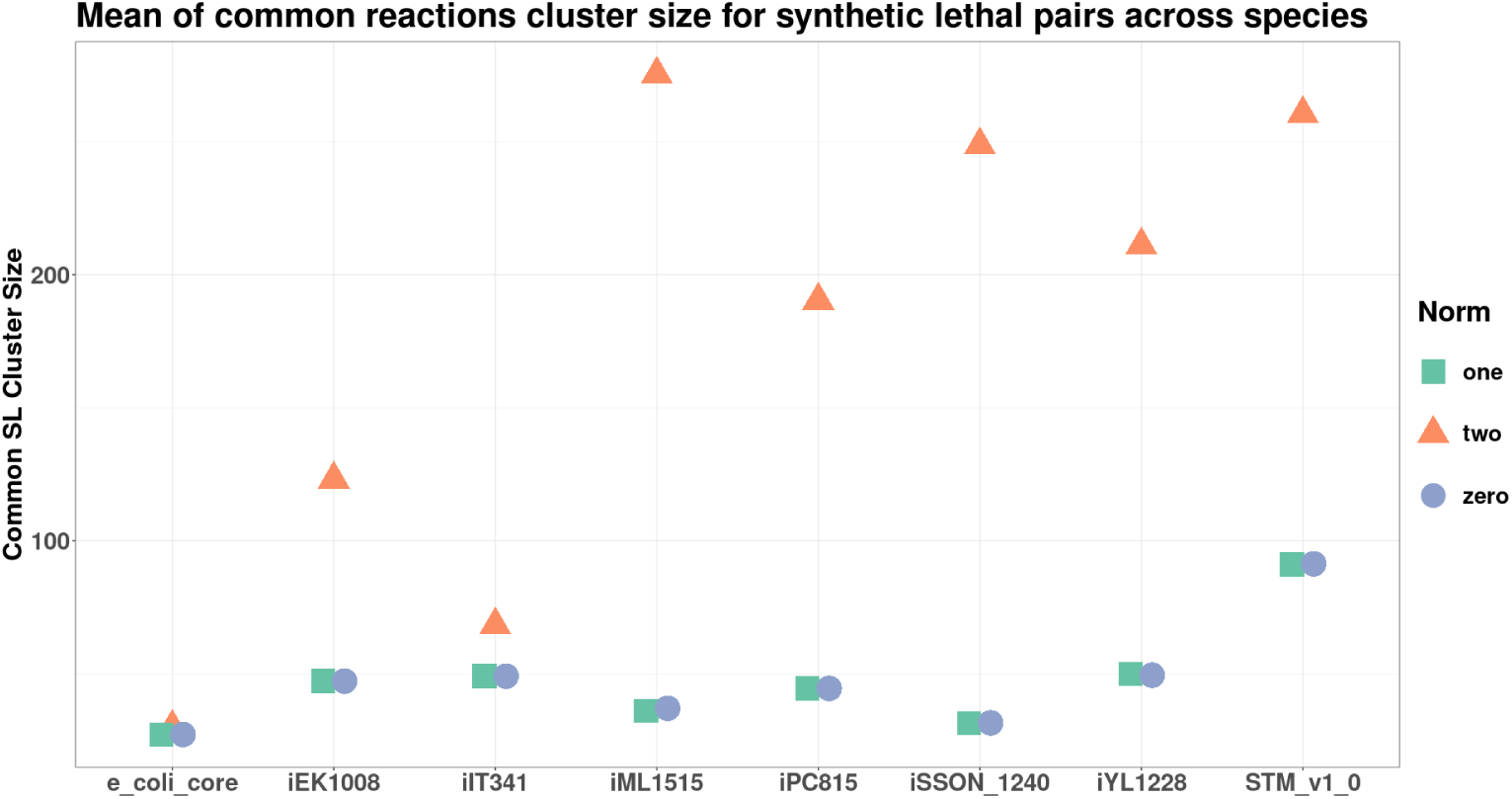
Distribution of Common SL Cluster Size across Organisms and Norms. Notice that the Common SL Cluster Size is the smallest for the zero and one norms, while the Common SL Cluster Size obtained using two norm is approximately an order of magnitude higher.

**Fig. A5.**
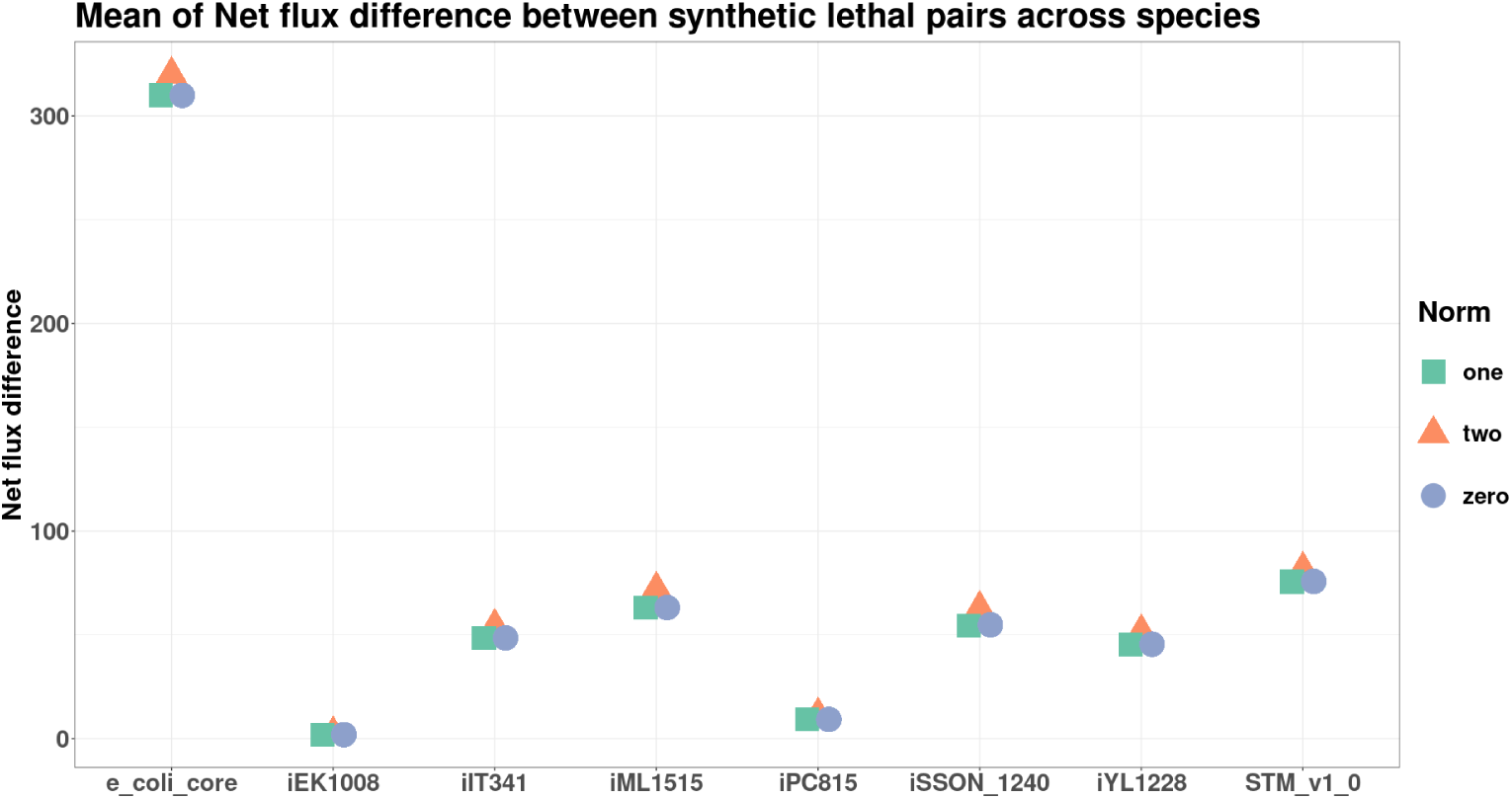
Distribution of Net flux difference between reaction pairs, across Organisms and Norms. The net flux difference shows similar results across the norms.

**Fig. A6.**
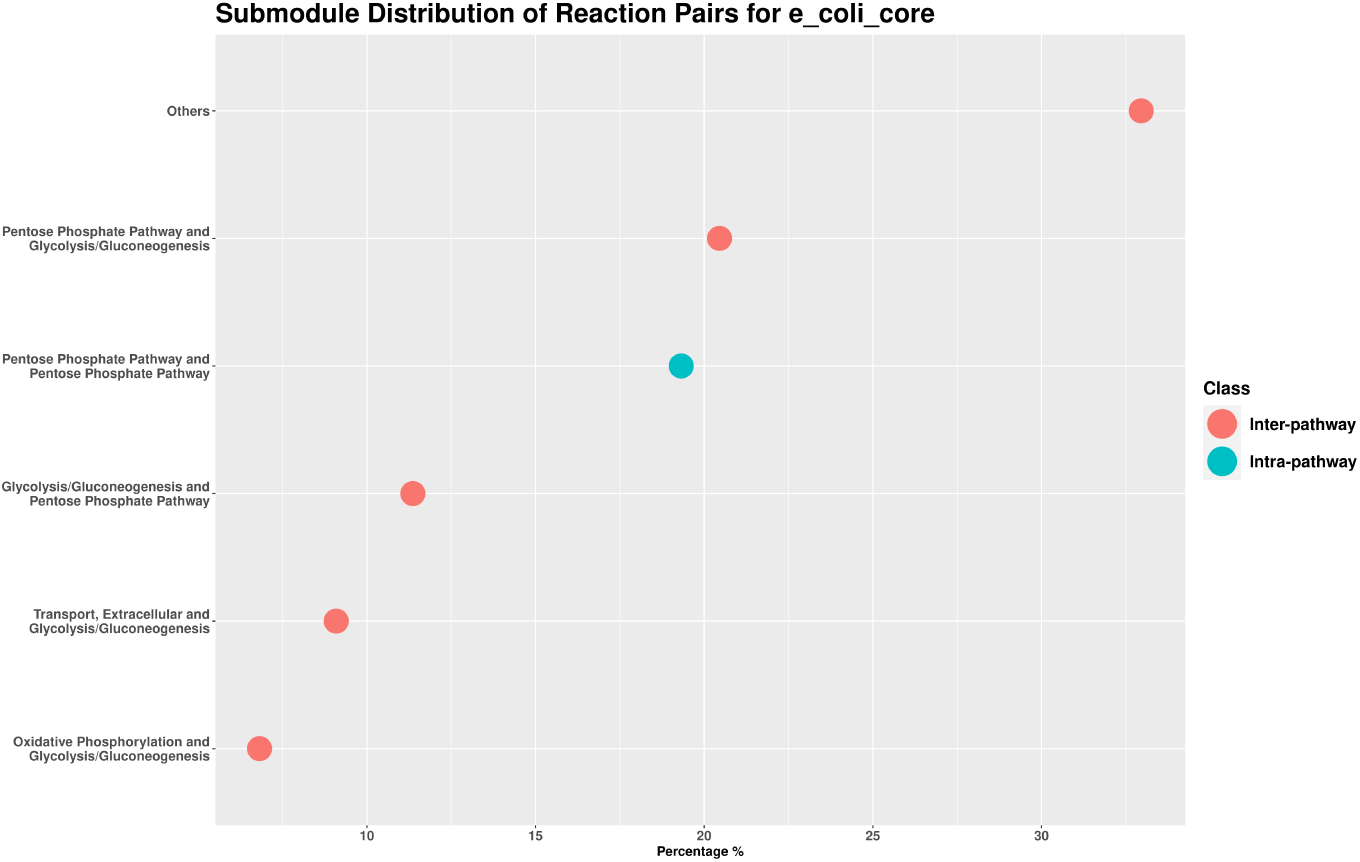
Metabolic Subsystem Analysis for the model e_coli_core. Distribution of DLs.

**Fig. A7.**
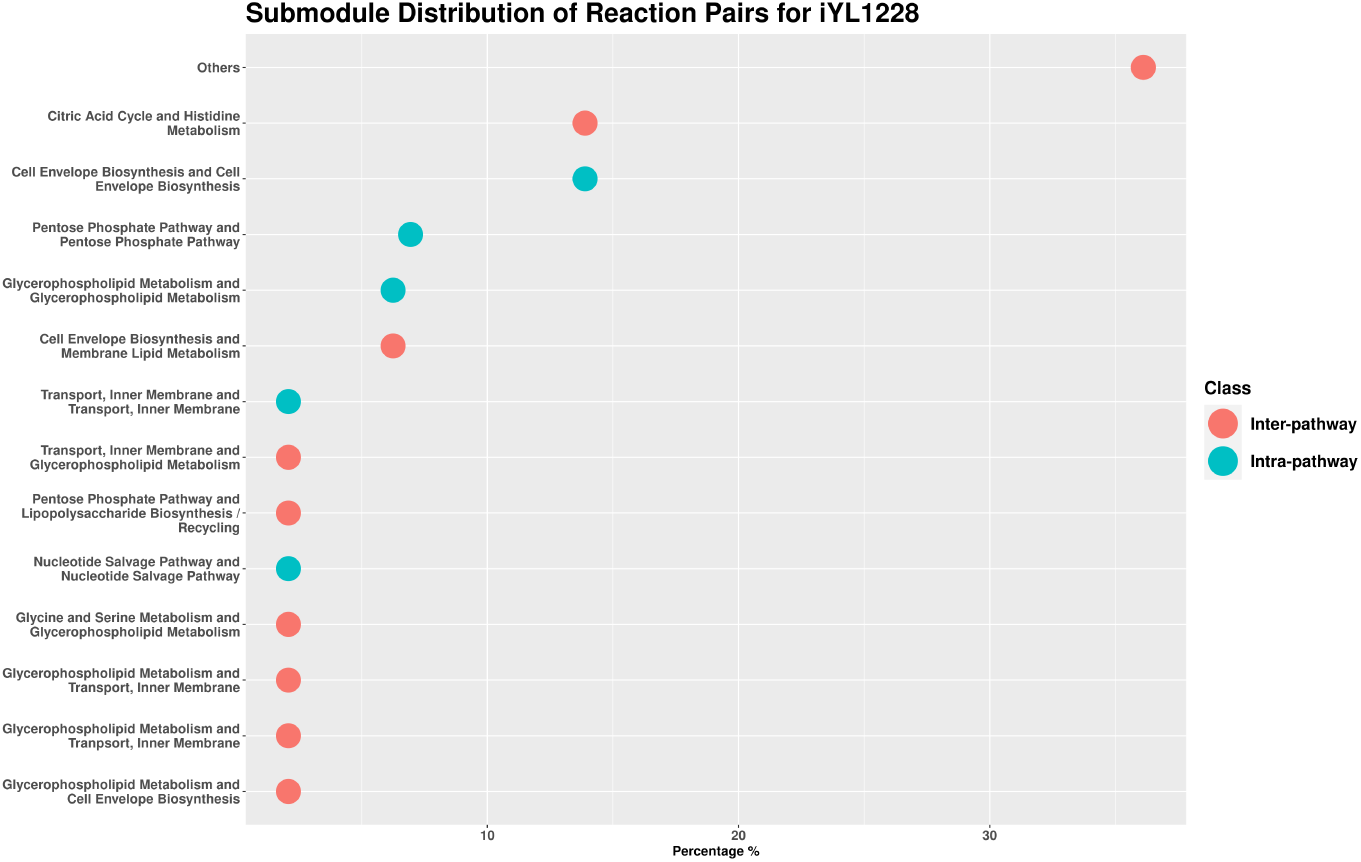
Metabolic Subsystem Analysis for the model *i*YL1228. Distribution of DLs.

**Fig. A8.**
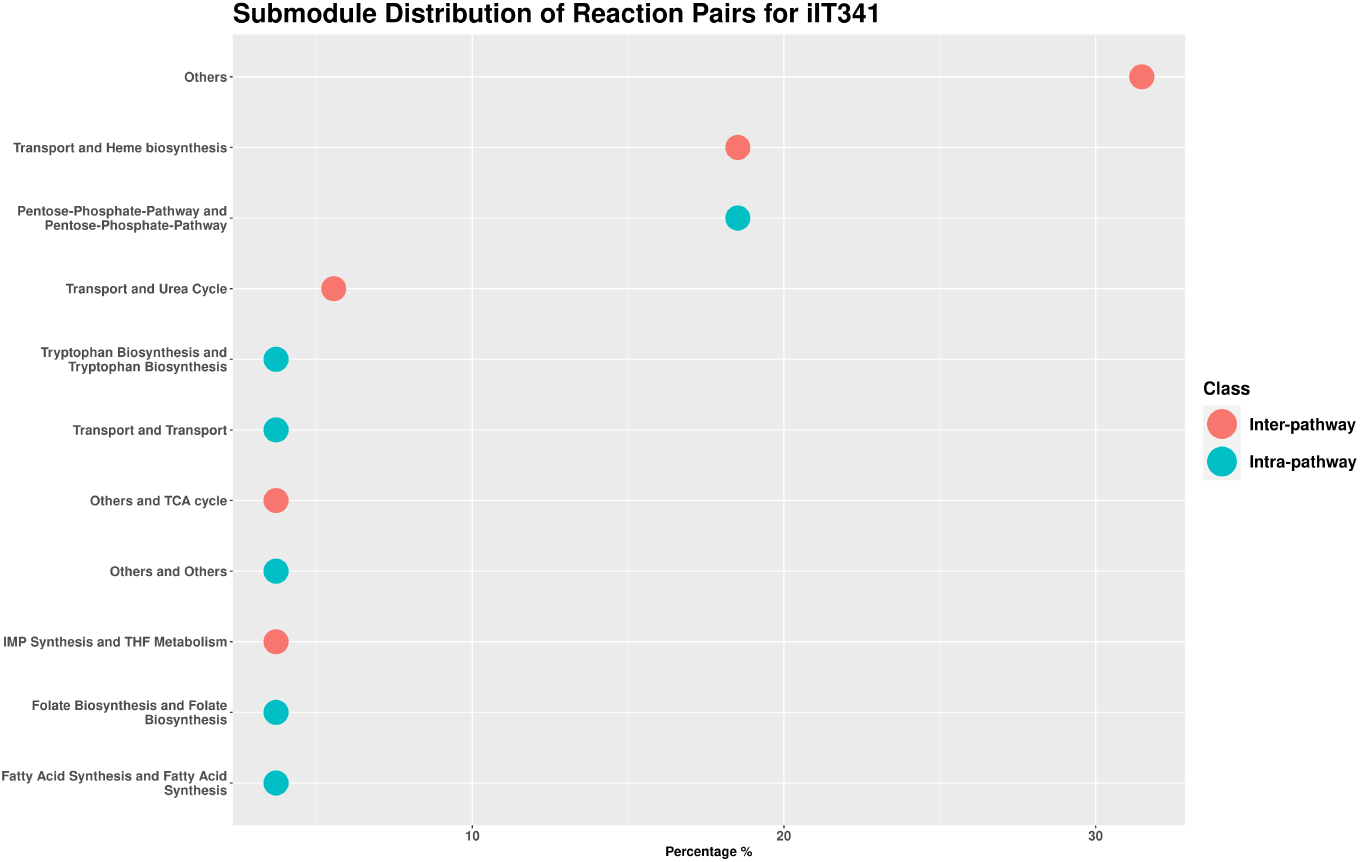
Metabolic Subsystem Analysis for the model *i*IT341. Distribution of DLs.

**Fig. A9.**
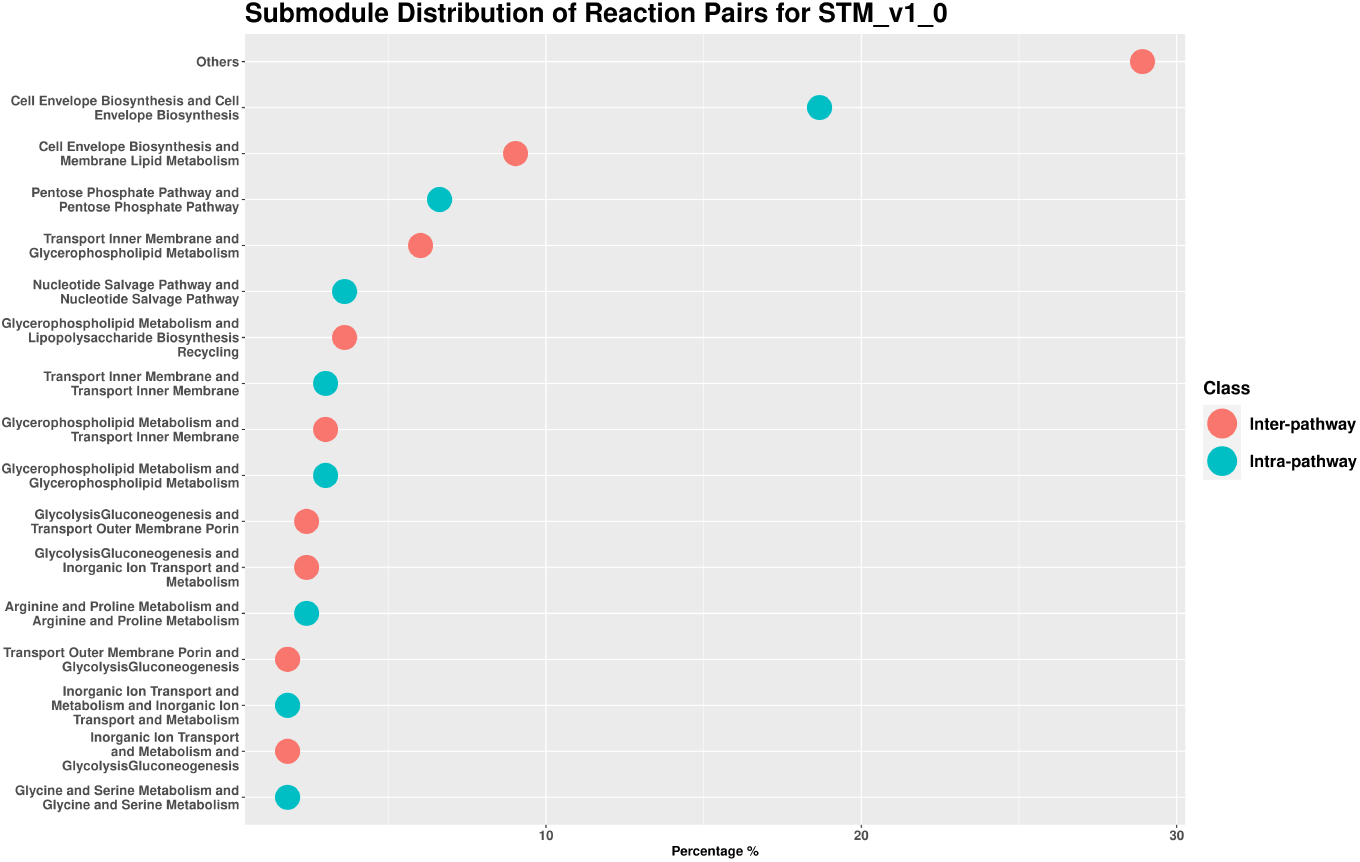
Metabolic Subsystem Analysis for the model STM_v1_0. Distribution of DLs.

**Fig. A10.**
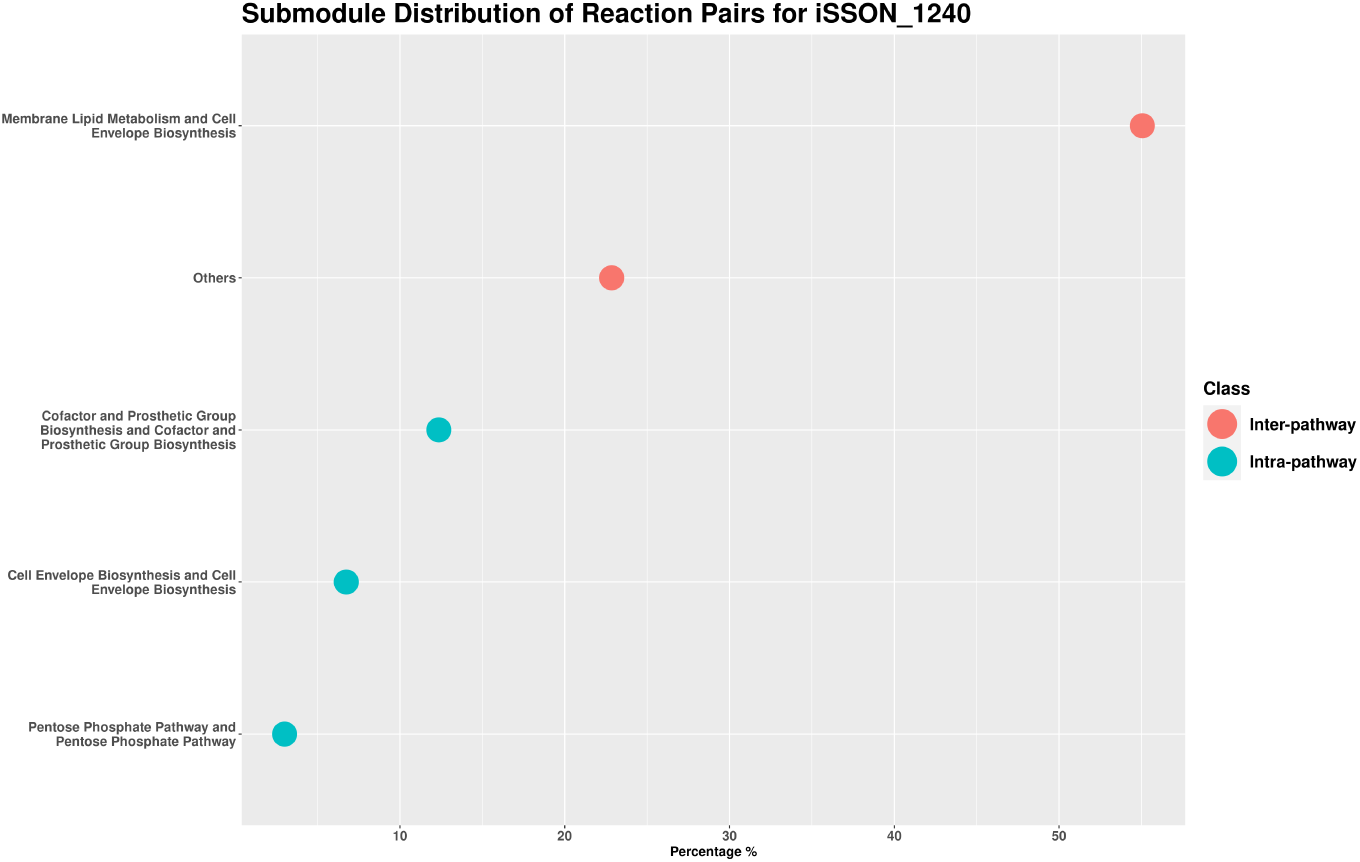
Metabolic Subsystem Analysis for the model *i*SSON_1240. Distribution of DLs.

**Fig. A11.**
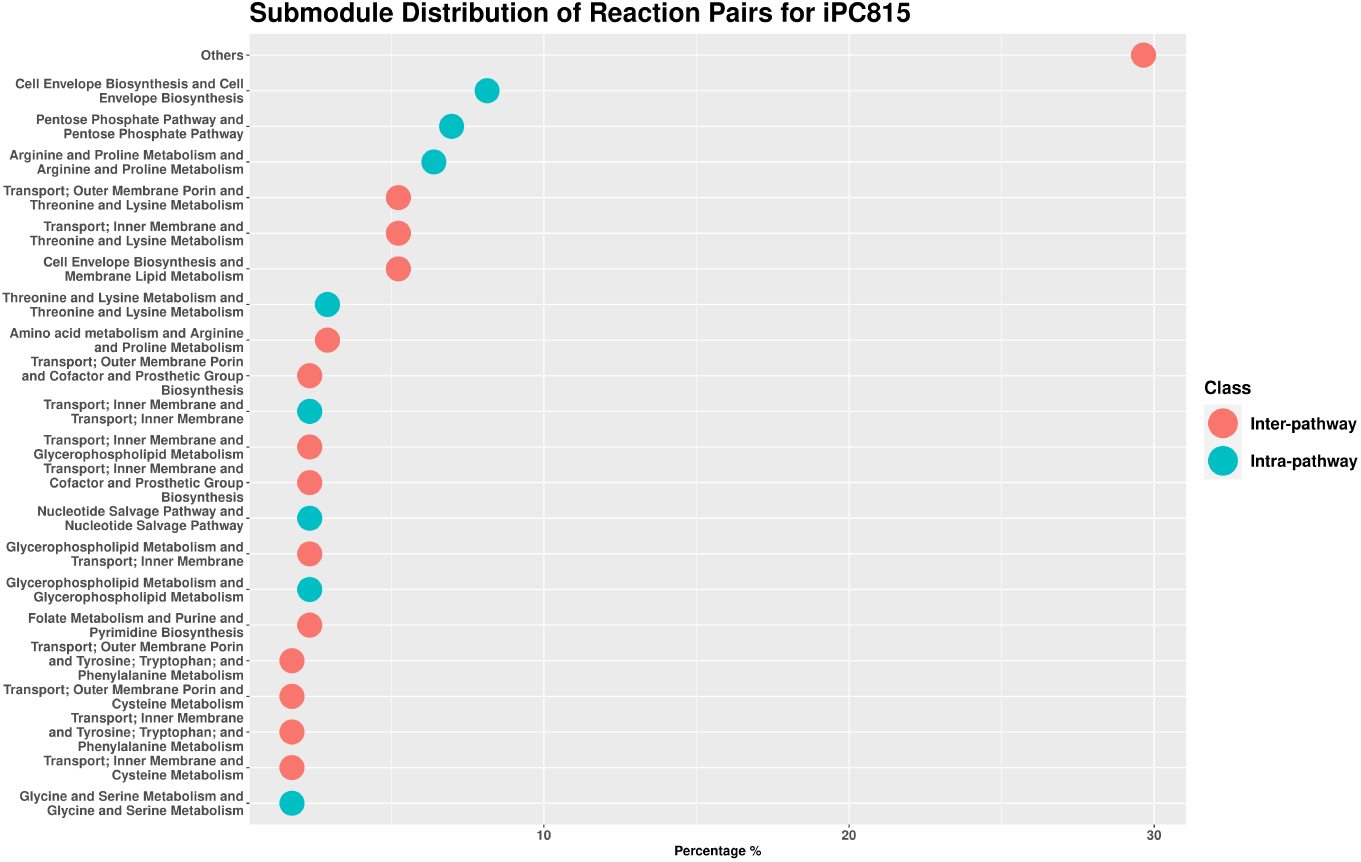
Metabolic Subsystem Analysis for the model iPC815. Distribution of DLs.

**Fig. A12.**
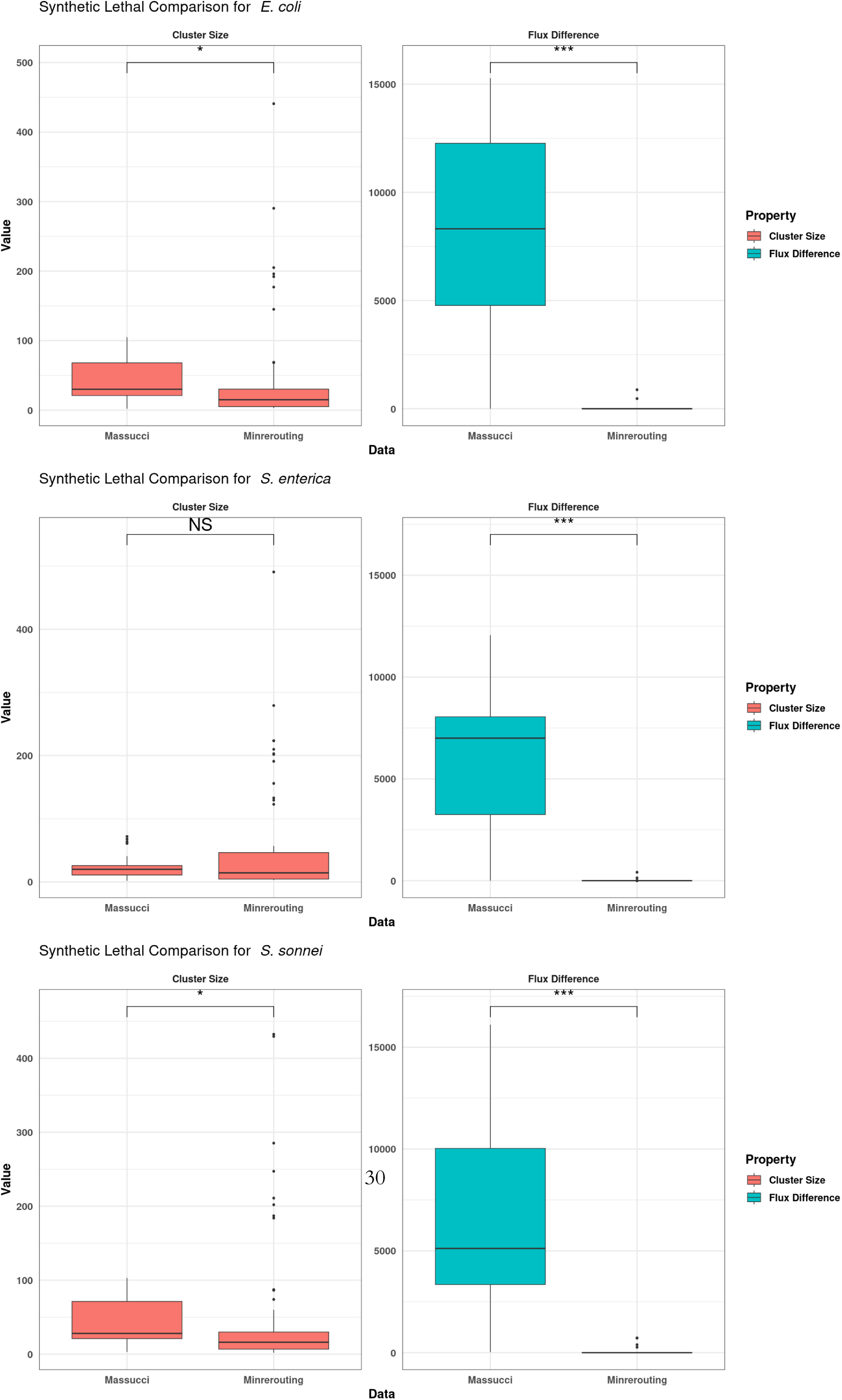
Comparison of Cluster Size and Net Flux Difference of Clusters for two different approaches to Synthetic Lethals—minRerouting and Massucci [17]. Notice that the net flux difference is significantly lesser for minRerouting than for the observations made previously. [17]. [Wilcoxon test was conducted. Significance levels: *** implies *p*-value < 0.001, ** implies *p*-value < 0.01, * implies *p*-value < 0.05]

**Fig. A13.**
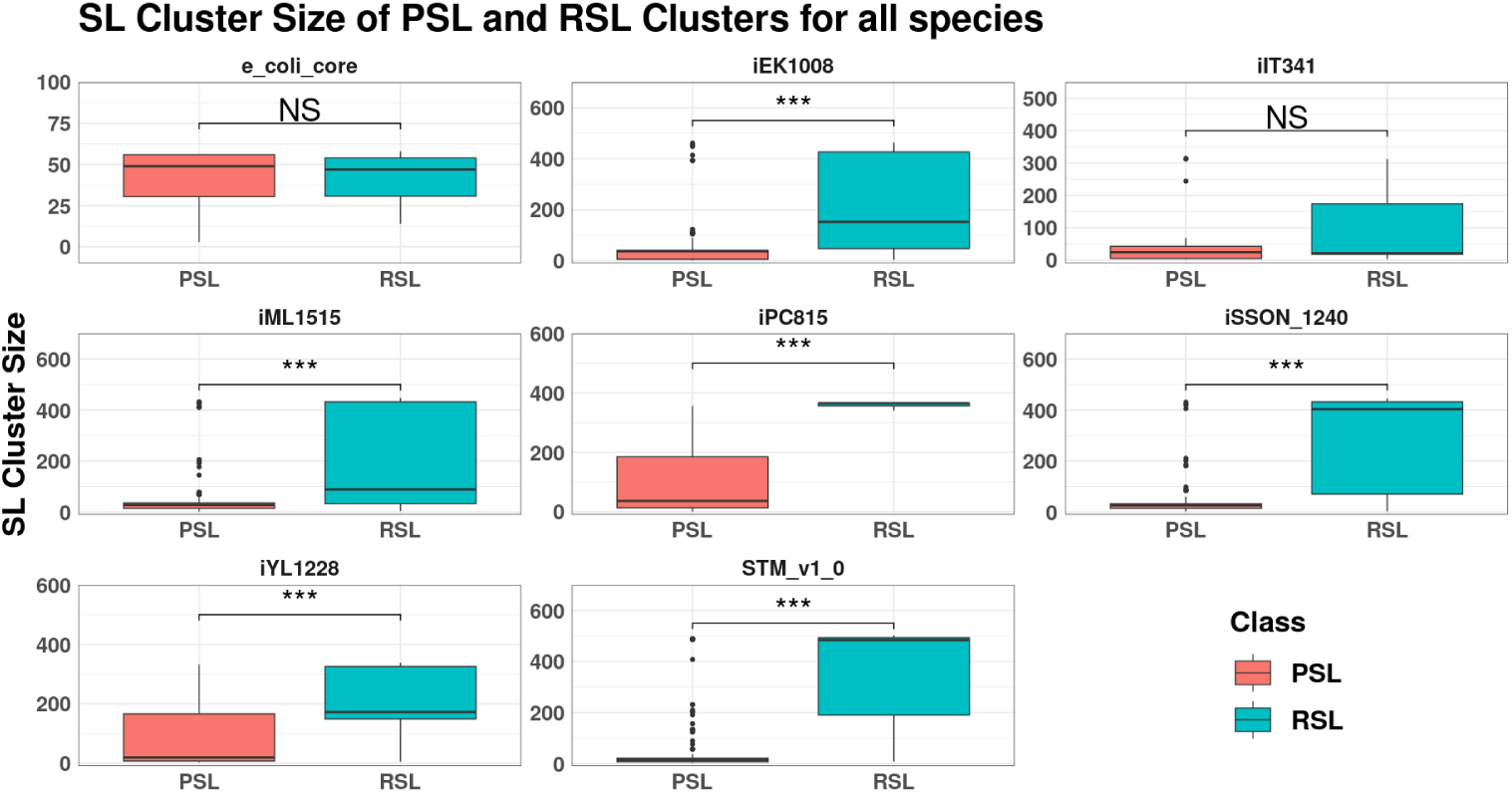
Distribution of Size of Clusters, across Organisms and Classes of the Cluster. The size is greater for RSLs than for PSLs. [Wilcoxon test was conducted. Significance levels: *** implies *p*-value < 0.001, ** implies *p*-value < 0.01, * implies *p*-value < 0.05]

**Fig. A14.**
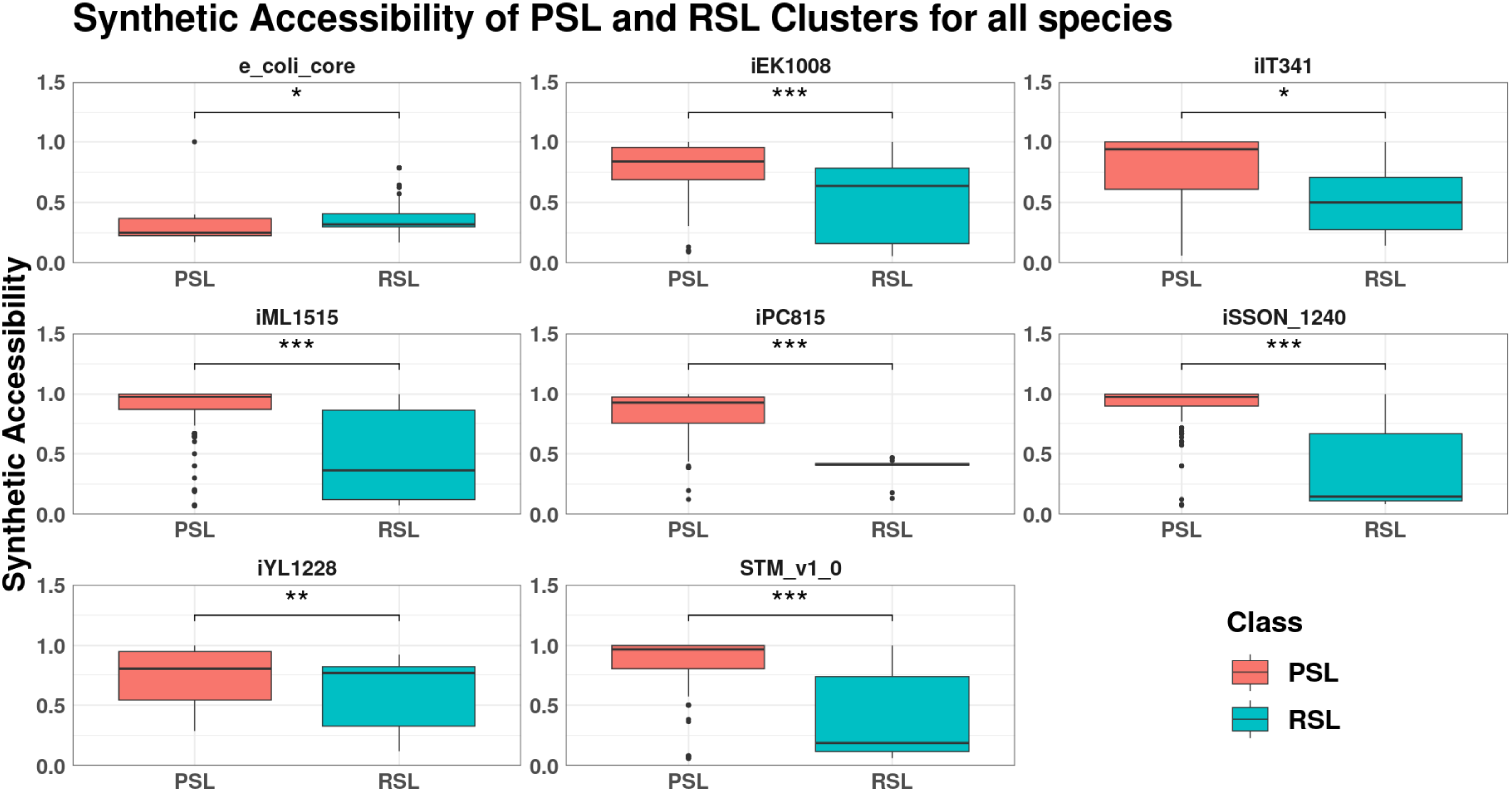
Distribution of Synthetic Accessibility of SL pairs, across Organisms and Classes of the Cluster. The Synthetic Accessibility is smaller for RSLs than for PSLs as PSLs activate new reactions while switching fluxes. [Wilcoxon test was conducted. Significance levels: *** implies *p*-value < 0.001, ** implies *p*-value < 0.01, * implies *p*-value < 0.05]

**Fig. A15.**
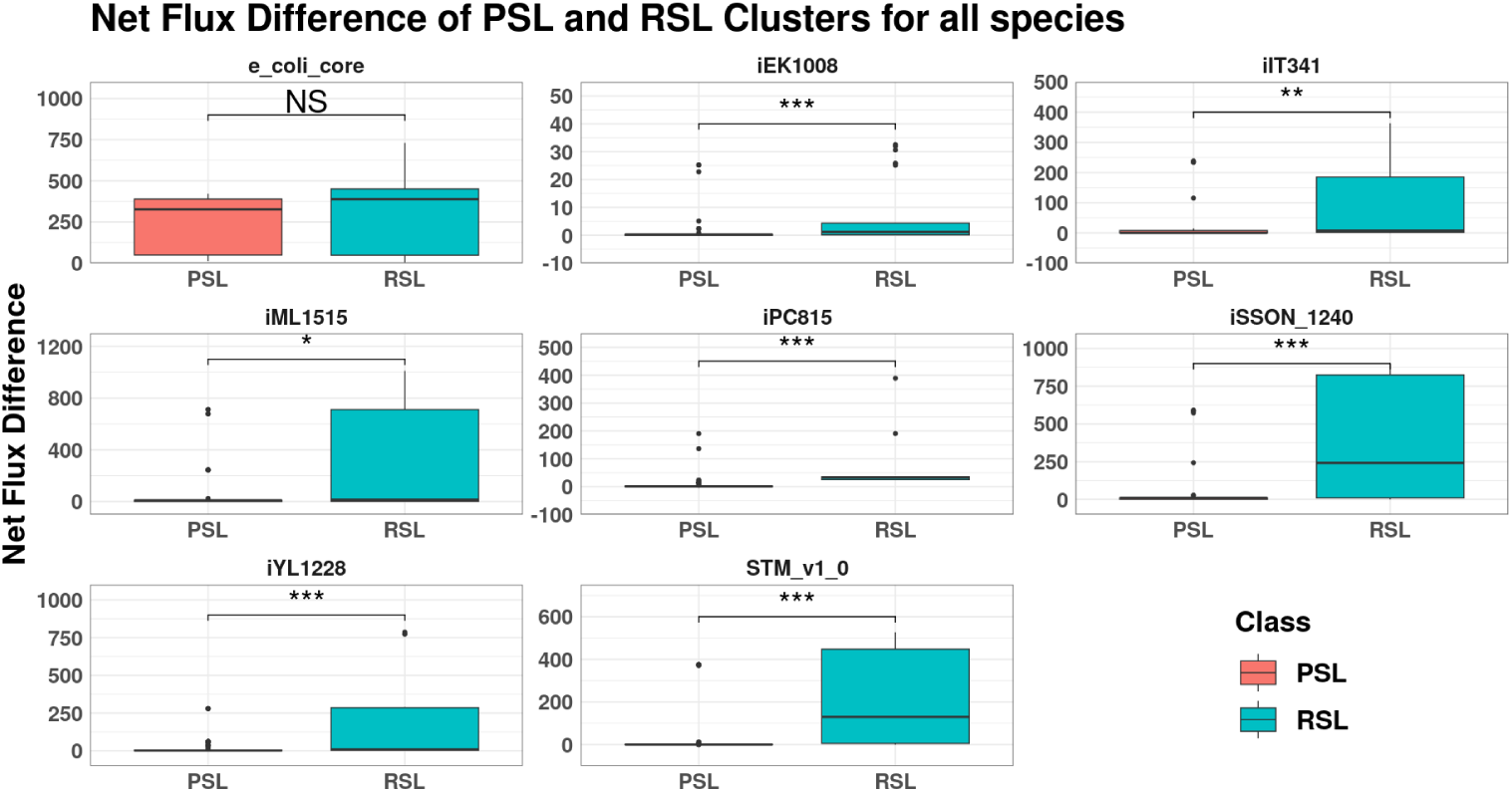
Distribution of Net Flux Difference of SL pairs, across Organisms and Classes of the Cluster. The net flux difference is also higher for RSLs than for PSLs. [Wilcoxon test was conducted. Significance levels-*** implies *p*-value < 0.001, ** implies *p*-value < 0.01, * implies *p*-value < 0.05]

**Fig. A16.**
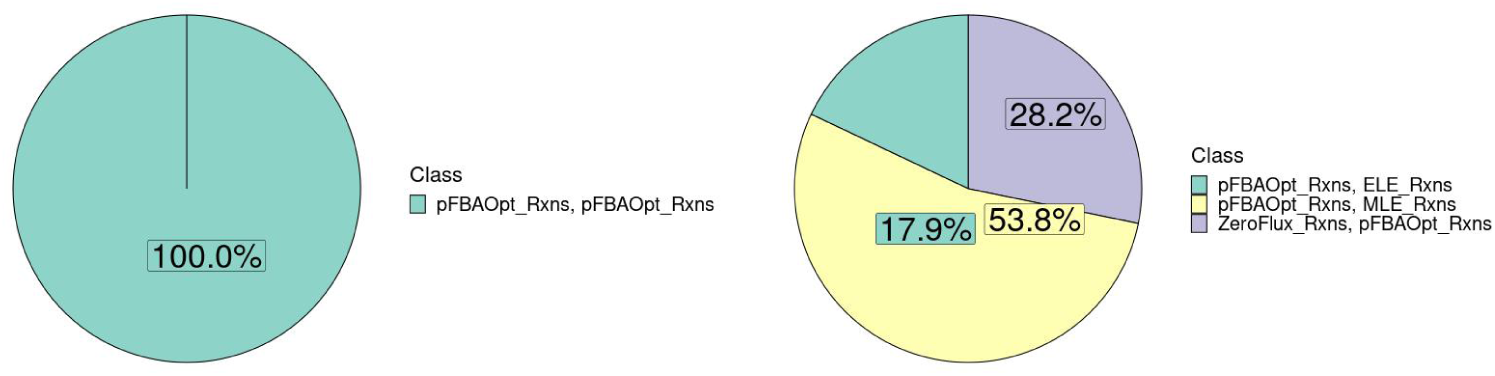
Schematic of reaction pair distribution between the RSL and PSL classes for *i*IT341.

**Fig. A17.**
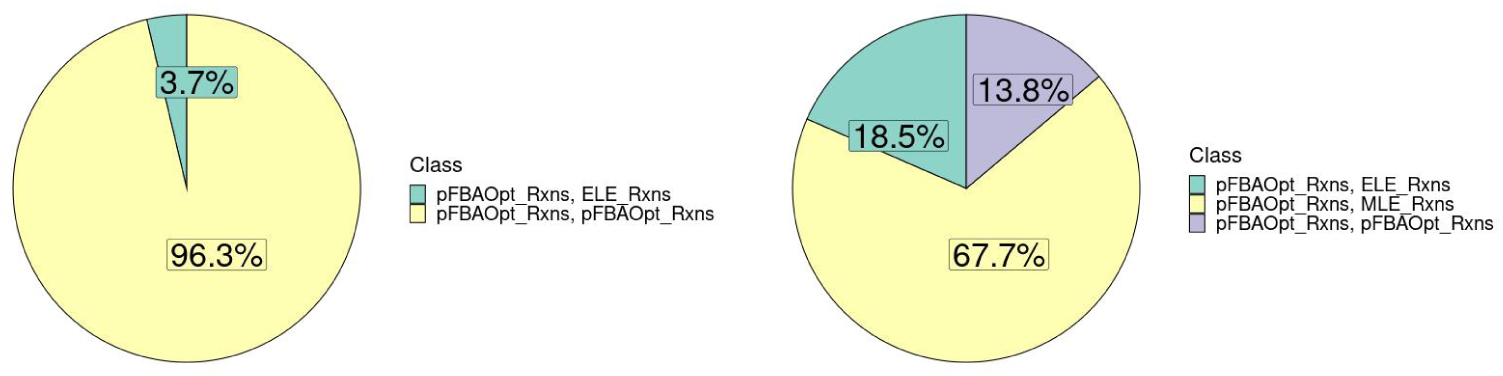
Schematic of reaction pair distribution between the RSL and PSL classes for *i*EK1008.

**Fig. A18.**
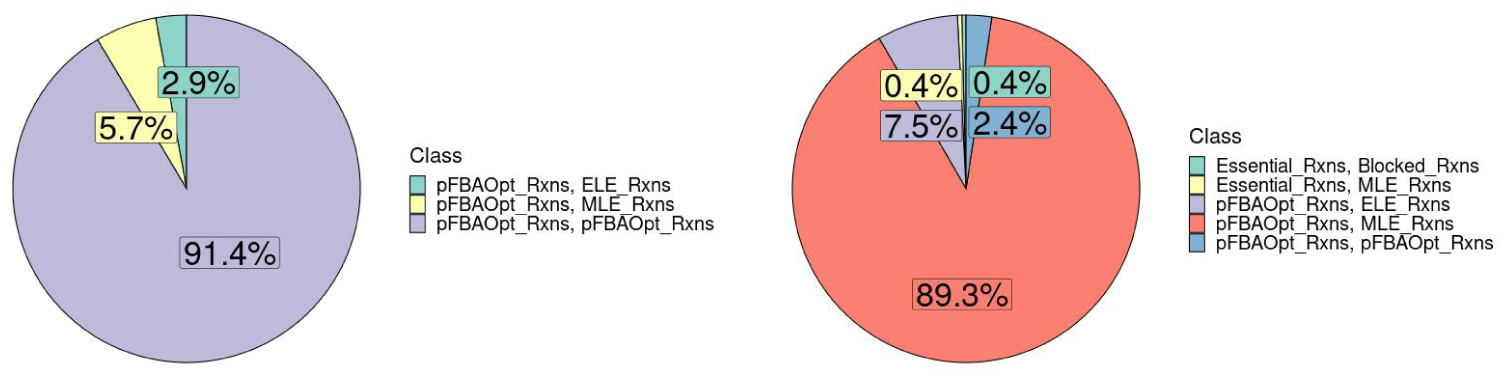
Schematic of reaction pair distribution between the RSL and PSL classes for *i*ML1515.

**Fig. A19.**
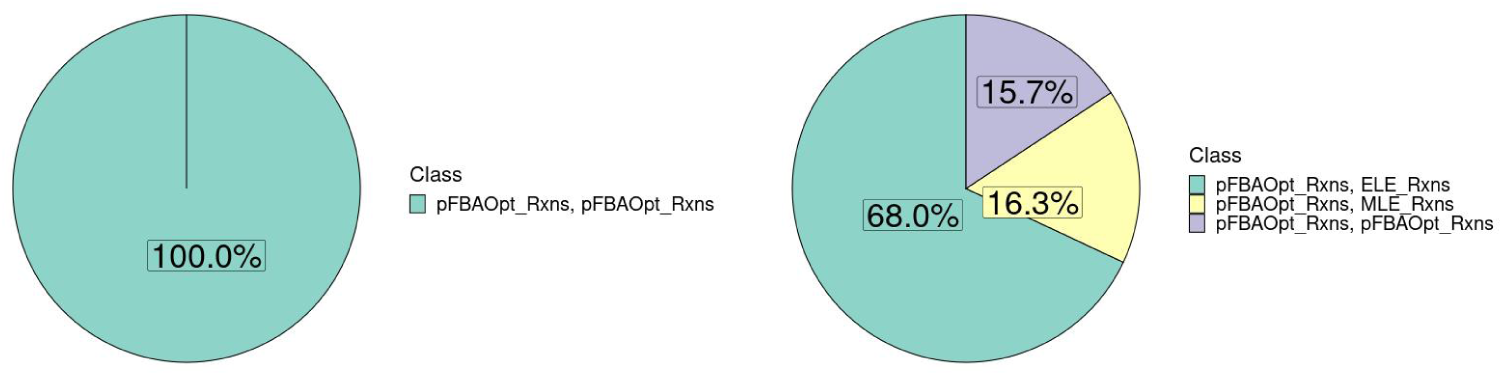
Schematic of reaction pair distribution between the RSL and PSL classes for *i*PC815.

**Fig. A20.**
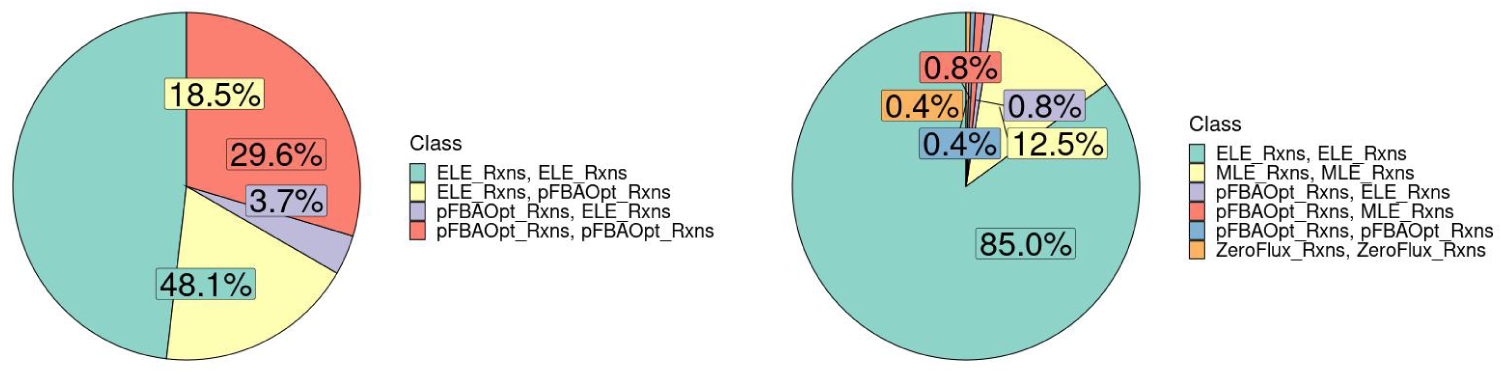
Schematic of reaction pair distribution between the RSL and PSL classes for *i*SSON_1240.

**Fig. A21.**
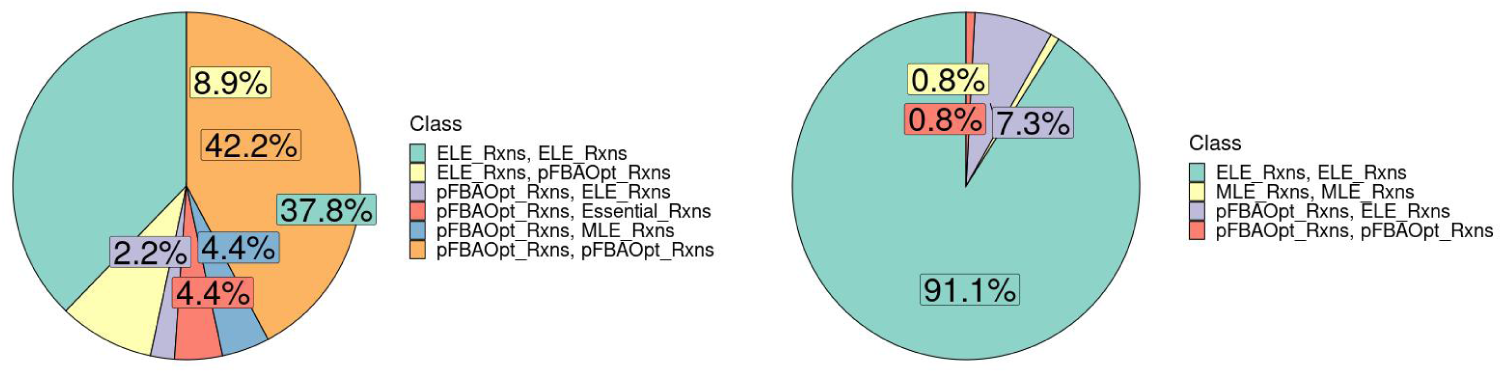
Schematic of reaction pair distribution between the RSL and PSL classes for STM_v1_0.

## Notes

http://bigg.ucsd.edu/

https://opencobra.github.io/cobratoolbox/stable/index.html

https://github.com/RamanLab/minRerouting

### Competing Interest Statement

The authors have declared no competing interest.

